# Metastatic dissemination of breast cancer stem cells requires MenaINV for lung extravasation but not survival

**DOI:** 10.64898/2026.01.21.700685

**Authors:** Mohd Nauman, Yookyung Jung, Burcu Ferrana Karadal, Suryansh Shukla, Camille L. Duran, Jiufeng Li, Prachiben Patel, Madeline Friedman-DeLuca, Nicole D. Barth, Robert Eddy, Wenjun Guo, John S. Condeelis, David Entenberg, Maja H. Oktay

## Abstract

Cancer stemness is a pivotal driver of tumor initiation, treatment resistance, and tumor cell survival. Cancer stem cells (CSCs), though constituting only a small fraction of primary tumor cells, are progressively enriched during metastatic progression: from circulating tumor cells traveling in the bloodstream, to disseminated tumor cells lodged in the lung vasculature, to extravasated tumor cells that have entered tissue parenchyma. However, whether CSCs have an intrinsic advantage for extravasation over cancer non-stem cells (CnSCs), or simply their increased representation in circulation renders them more likely to extravasate, remains unresolved. MenaINV, an invasive isoform of the actin regulatory protein Mena, promotes tumor cell transendothelial migration in primary and secondary sites, yet the direct mechanistic link between stemness and MenaINV in lung metastasis remains unresolved. Here, using a validated fluorescent stemness reporter (SORE6) to identify CSCs, we found that CSCs display elevated MenaINV expression relative to CnSCs. High-resolution intravital imaging showed that CSCs extravasate efficiently into lung parenchyma and survive at higher levels, robustly forming metastatic lesions, while CnSCs show limited extravasation, low survival, and poor colonization. Mechanistically, MenaINV disruption in CSCs specifically impaired extravasation without affecting survival, demonstrating that MenaINV is the key extravasation effector downstream of stemness, whereas stemness-associated factors independently confer survival advantages. Moreover, reintroduction of MenaINV in CnSCs restores their extravasation ability upon which extravasated CnSCs reactivate stem program and form metastases. Overall, we discovered a hierarchical framework where stemness regulates both survival and extravasation capacity, with MenaINV as the key CSC extravasation effector.

Significance: This study reveals how breast cancer stem cells achieve metastatic dominance through separable pathways: MenaINV-dependent extravasation and MenaINV-independent survival, providing rationale for targeting stem program to improve patient outcome.

## Introduction

Breast cancer is the most common cancer among women and the second leading cause of cancer-related mortality in the United States (Giaquinto, Sung et al. 2024). The majority of cancer deaths are attributed to metastatic disease (Dillekas, Rogers et al. 2019), a process in which tumor cells disseminate from primary tumors, survive in circulation, extravasate at secondary sites such as the lungs, and form metastatic foci in these distant organs (Mehlen and Puisieux 2006, DeSantis, Ma et al. 2019, Borriello, Condeelis et al. 2021). During this cascade, circulating tumor cells (CTCs) travel through the bloodstream until they become physically lodged within the microvasculature of distant organs such as the lung, at which point they are termed disseminated tumor cells (DTCs). These intravascular DTCs must then extravasate across the endothelial barrier to enter tissue parenchyma and establish metastatic lesions. Numerous studies have demonstrated that cancer stem cells (CSCs), a subpopulation characterized by enhanced tumor initiation capacity, treatment resistance, and superior tumor cell survival, play a critical role in primary tumor formation (Shackleton, Quintana et al. 2009, Nassar and Blanpain 2016, Zhao 2016, Papadaki, Stoupis et al. 2019, Sharma, Tang et al. 2021) but their role in metastasis formation remains less well elucidated (Amey and Karnoub 2017, Czarnogorski, Czernicka et al. 2025).

We have previously shown that CSCs constitute only a minor subpopulation within primary tumors, but that their proportion dramatically increases during metastasis. We found CSCs to be enriched in close proximity to TMEM doorways, portals within the primary tumor vasculature where cancer cells intravasate (Sharma, Tang et al. 2021). Furthermore, our intravital imaging of primary tumors showed that CSCs display a slow migratory and invadopodium-rich phenotype indicative of disseminating tumor cells that are capable of degrading vascular basement membranes (Gligorijevic, Bergman et al. 2014, Sharma, Tang et al. 2021). Consistently, we observed that CSCs comprise approximately 66% of circulating tumor cells, 77% of single cells found both intravascularly and extravascularly in the lung, and 84% of cells in early metastatic lesions (Sharma, Tang et al. 2021). This high percentage of CSCs that successfully complete the initial steps of the metastatic cascade raises the question of whether CSCs possess inherent mechanisms that give them extravasation and survival advantages over cancer non-stem cells (CnSCs), or if this observation is simply a consequence of their increased prevalence in circulation.

One of the mechanisms by which tumor cells are able to intravasate in primary tumors (Roussos, Goswami et al. 2011, Pignatelli, Goswami et al. 2014) and extravasate at secondary sites (Borriello, Coste et al. 2022) is through the expression of MenaINV, an invasive isoform of the actin regulatory protein Mena (Gertler and Condeelis 2011). MenaINV has been shown to convey the slow migratory invadopodium-rich phenotype in disseminating cancer cells (Roussos, Balsamo et al. 2011, Pignatelli, Bravo-Cordero et al. 2016, Sharma, Tang et al. 2021) and we found MenaINV-expressing cells are also enriched near TMEM doorways (Sharma, Tang et al. 2021, Borriello, Coste et al. 2022).

Together these findings raise the possibility that MenaINV might act as a critical molecular effector enabling CSCs to accomplish key steps in metastasis, such as efficient transendothelial migration and survival at secondary sites (by escaping the harsh confines of the vasculature faster). Supporting this conjecture is our finding that over 50% of CSCs in breast tumors co-express MenaINV (Sharma, Tang et al. 2021), suggesting the cooperative role of stemness and MenaINV in intravasation, and possibly in subsequent extravasation. Additionally, published literature demonstrates that genes and pathways which regulate stemness (such as Sox2, Oct4, and Sox9), are also regulators of EMT (Wang, Lu et al. 2014, Liu and Guo 2021).

We therefore hypothesized that stemness and MenaINV work together in facilitating tumor cell extravasation and survival in the lung. To address this hypothesis, we used an experimental metastasis model in which equal numbers of CSCs and CnSCs were introduced into the circulation. CSCs and CnSCs were identified by our validated SORE6 reporter that marks stem cells based on their expression of Sox2 and Oct4 (Tang, Raviv et al. 2015, Sharma, Tang et al. 2021), core transcription factors that regulate the stem cell state (Chu, Tian et al. 2024). Intravital multiphoton imaging of the live murine lung was then performed to track the fate of individual disseminated tumor cells. In the current study, we use more specific terminology than our prior work (Sharma, Tang et al. 2021, Borriello, Coste et al. 2022) to distinguish tumor cell populations based on their precise anatomical location. We define intravascular cells as tumor cells physically lodged within the lung microvasculature and extravasated cells as those that have crossed the endothelial barrier into the lung parenchyma. This distinction allows us to separately assess the extravasation and survival capacities of CSCs versus CnSCs. This approach allowed us to directly measure and compare the extravasation and survival of CSCs vs CnSCs in the lung microenvironment, as well as to determine the contribution of MenaINV to these processes.

## Results

To test the hypothesis that CSCs intrinsically differ from CnSCs in their ability to extravasate and survive within the lung microenvironment, we employed time-lapse intravital multiphoton imaging in conjunction with our validated SORE6 reporter system to directly compare these cell populations following their introduction into the circulation. This approach enables dynamic, high-resolution visualization and fate tracking of CSCs and CnSCs in the living lung, permitting quantitative assessment of their rates of extravasation and subsequent survival. By continuously monitoring these distinct populations in situ, we generated real-time evidence to address whether CSCs possess a functional advantage during the early steps of lung colonization. As a first step in this approach, we established a cell line containing our previously published reporter for stemness. The SORE6 reporter has been validated to accurately identify cancer stem cells using several methods, including serial dilution and in vivo tumor-initiation assays, as well as confirmation of the enrichment of canonical stem cell markers and transcription factors in the SORE6-positive population relative to SORE6-negative cells, (Tang, Raviv et al. 2015, Sharma, Tang et al. 2021).

The SORE6 reporter construct contains six tandem repeats of a composite Sox2/Oct4 response element coupled to a minimal cytomegalovirus promoter (minCMV) (**Fig. 2A and Supplementary Fig. 1A**). Upon activation by the CSC master transcription factors Sox2 and Oct4, this promoter drives expression of a destabilized fluorescent protein (dsCopGFP), a C-terminally truncated CD19 marker (for flow-based identification of cells that have successfully incorporated the construct), and a FLAG epitope tag. For live cell imaging, stemness is detected via GFP fluorescence, while in fixed tissues stemness is detected via FLAG immunostaining, as dsCopGFP degrades upon fixation. dsCopGFP was chosen over GFP for in vivo work as the destabilized fluorescent protein’s rapid turnover results in a more temporally responsive indicator of stemness-associated transcriptional activity (Sharma, Tang et al. 2021).

Throughout this manuscript, we use simplified terminology to distinguish cell populations. All experimental cells constitutively express tdTomato as a volume marker. For clarity, we designate tdTomato+CD19+minCMV control cells (lacking Sox2/Oct4 response elements) as minCMV control cells, tdTomato+CD19+SORE6-cells (GFP fluorescence below threshold) as CnSCs or SORE6-cells, and tdTomato+CD19+SORE6+ cells (GFP fluorescence above threshold) as CSCs or SORE6+ cells.

We transduced MDA-MB-231-tdTomato cells with the SORE6 reporter lentivirus. Using the minCMV control to establish the GFP threshold, we flow sorted cells into SORE6-(CnSC) and SORE6+ (CSC) populations based on GFP fluorescence (**Supplementary Fig. 2**). GFP expression in sorted populations was confirmed by fluorescence imaging (**Supplementary Fig. 3A**).

### Expression of stem cell markers is elevated in SORE6+ compared to SORE6-cell populations

To verify that sorting based upon the SORE6 reporter fluorescence indeed identified cells with increased Sox2/Oct4 transcription we flow sorted cells taken from culture into SORE6- and SORE6+ populations based upon their GFP expression and determined their expression levels using quantitative RT-PCR (qRT-PCR). We observed significantly higher expression of both Sox2 and Oct4, as well as several other established markers of cancer stemness in SORE6+ compared to SORE6-cells (**Supplementary Fig. 3B**).

These findings confirmed that the reporter functioned as intended, and, together with previous studies validating SORE6+ cells as cancer stem cells (Sharma, Tang et al. 2021), supported the use of SORE6 reporter activity as a reliable indicator of cancer stemness in our cells. Based on this evidence, we hereafter designate SORE6-cells as CnSCs and SORE6+ cells as CSCs.

### Breast cancer stem cells (CSCs) have elevated expression of MenaINV compared to cancer non-stem cells (CnSCs), both in vitro and in vivo

We previously found that over 50% of CSCs in primary breast tumors co-express MenaINV (Sharma, Tang et al. 2021). Thus, to investigate the connection between stemness and MenaINV expression, we measured the mRNA levels of MenaINV, Mena11a, and pan-Mena (which detects all Mena isoforms) in CnSCs and CSCs. We observed that, compared to CnSCs, CSCs exhibit significantly higher MenaINV expression, while the levels of Mena11a and pan-Mena do not differ between CnSCs and CSCs (**Fig. 1A**). Immunofluorescence staining further confirmed that, in vitro, CSCs on average express higher MenaINV protein levels compared to CnSCs by microscopy (**Fig. 1B & 1C**). We also checked the correlation between stemness and MenaINV in individual cells in vivo in primary orthotopic tumors generated via injection of SORE6 reporter-containing cells and co-staining harvested tissues with MenaINV and FLAG antibodies (**Fig. 1E**) and observed a strong correlation between SORE6 reporter expression (via FLAG) and MenaINV (**Fig. 1F**).

**Figure 1.**
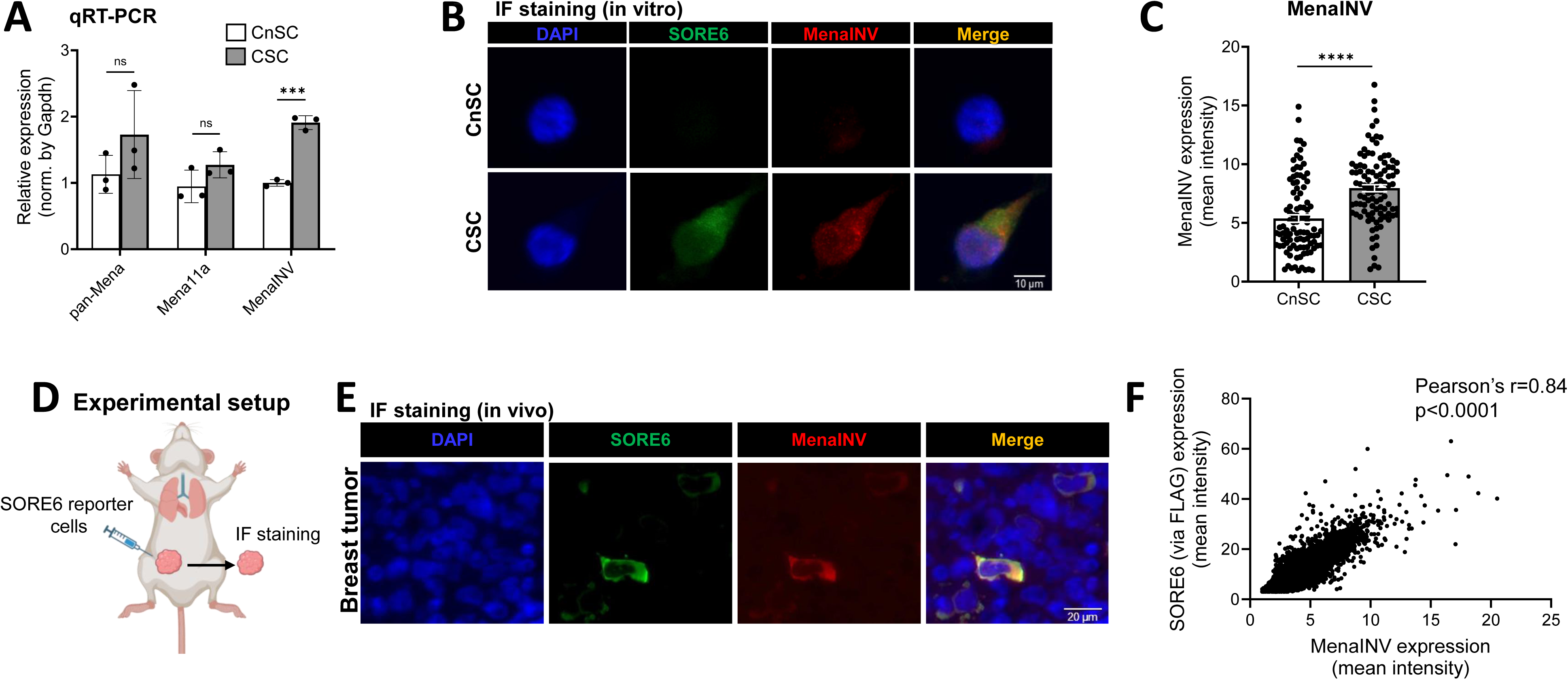
Breast cancer stem cells (CSCs) have elevated expression of MenaINV in vitro and in vivo. **A)** Quantitative RT-PCR analysis showing that higher MenaINV expression in FACS-sorted cancer stem cells (CSCs) compared to cancer non-stem cells (CnSCs), while no difference is observed for Mena11a and pan-Mena. Error bars = mean ± SEM (n=3 independent assays), normalized to GAPDH and quantified using the log2ddCT method. Student’s t-test, ***p<0.0002. **B)** Representative immunofluorescence images demonstrating SORE6 reporter (GFP) and MenaINV staining in CnSCs and CSCs. **C)** Quantification of MenaINV protein expression per cell in CnSCs and CSCs (n = 100 cells/group, pooled from n = 3 independent wells/group). Dots represent individual cells. Mann–Whitney U test, ****p<0.001. **D)** Schematic of experimental setup for orthotopic generation of primary breast tumors created with Biorender. **E)** Representative fluorescence image showing co-localization of SORE6 reporter (detected via FLAG) and MenaINV in primary breast tumor tissue. **F)** Correlation analysis of mean FLAG and MenaINV intensity across all cells detected in n=3 tumors. Abbreviations: CSCs, cancer stem cells; CnSCs, cancer non-stem cells; SEM, standard error of the mean.

### CSCs and CnSCs do not appreciably interconvert on the minutes-to-hours time frame

Our previous work demonstrated that ∼66% of circulating tumor cells (CTCs) in mice bearing primary breast tumors were CSCs and that this percentage increased further among disseminated tumor cells (DTCs) lodged in the lung vasculature, with ∼77% identified as CSCs (Sharma, Tang et al. 2021). These findings raise the question of whether the enrichment of CSCs in lung metastases reflects intrinsic advantages in extravasation and survival, or simply their increased representation in circulation. To understand the dynamics of CSCs and CnSCs, we injected equal ratios of these cells retro-orbitally into immunocompromised mice bearing implantable lung imaging windows.

In our experimental setup, MDA-MB-231 cancer cells are transduced with the SORE6 reporter construct (**Fig. 2A)** and tdTomato (and additionally labeled with a red cell tracker), were flow sorted into separate SORE6- and SORE6+ populations, and mixed together at a 1:1 ratio (**Fig. 2B**). Mice bearing lung imaging windows were retro-orbitally injected with a far-red dextran (to visualize the blood vasculature) as well as the mixed cancer cells (**Fig. 2C**) and imaged over time to determine their stem status. Cells which expressed GFP fluorescence above the minCMV level were considered CSCs (**Supplementary Fig. 4**). We performed continuous time-lapsed intravital imaging (minutes frame rate) of cells over time periods from 0-8, 16-24, and 24-30, 48-51, and 56-59 hours. Consistent with our prior observations of disseminated tumor cells in the lung, motility was minimal during this period (**Supplementary Fig. 5 and Supplementary Movie 1**). Notably, throughout our observation, very few CSCs and CnSCs (3 of 79 cells = 3%) were observed to undergo interconversion over 60 hours. Instead, their respective identities remained stable and distinct within the lung microenvironment. These findings indicate a lack of plasticity between CSC and CnSC states during the critical initial phase after arrival to the lung, suggesting that the enhanced metastatic potential of CSCs is maintained by a stable cell identity rather than dynamic conversion between stem and non-stem states.

**Figure 2.**
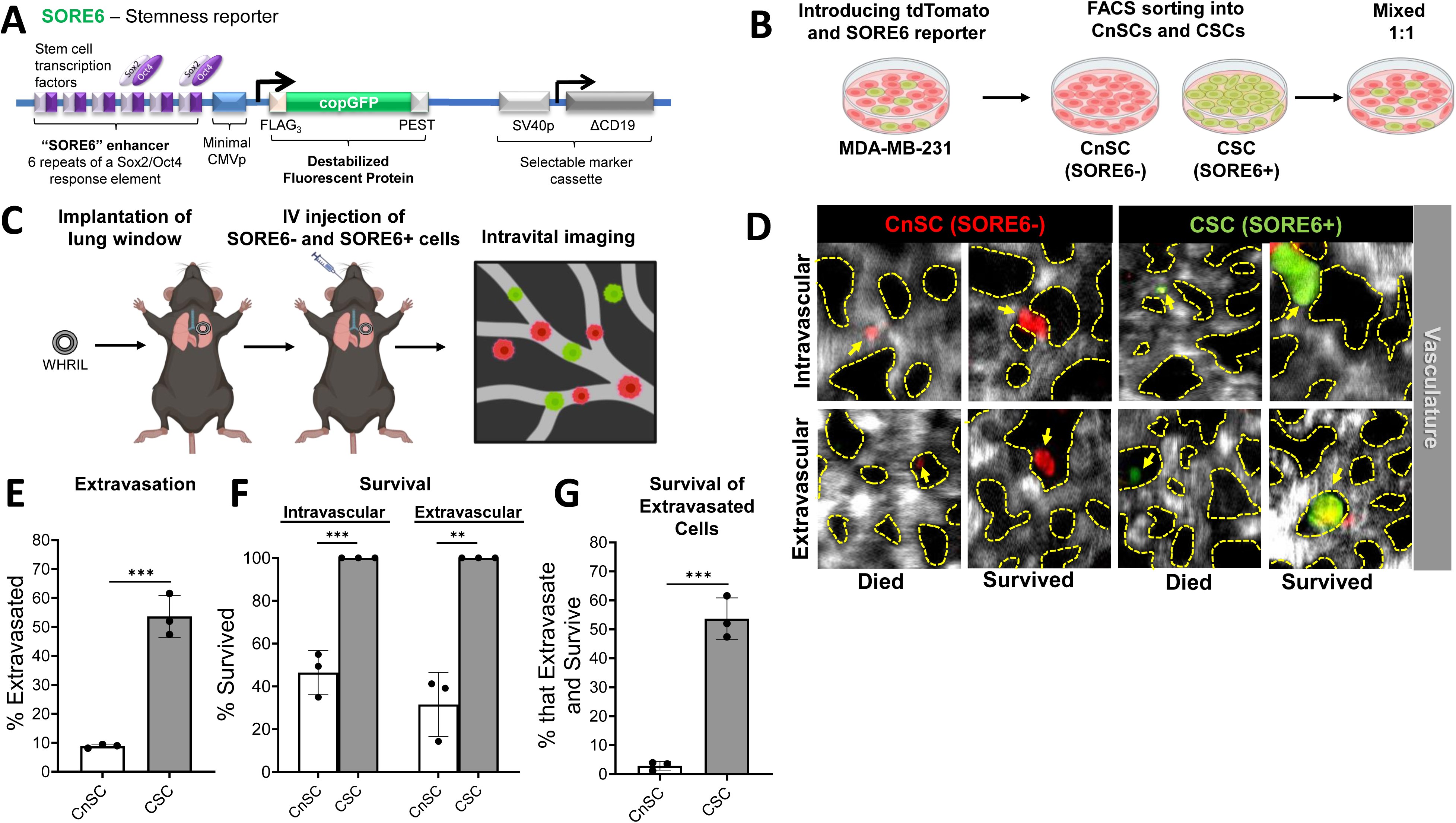
Intravital imaging reveals more efficient extravasation and better survival of cancer stem cells compared to cancer non-stem cells. **A)** Schematic of SORE6 reporter construct. **B)** MDA-MB-231 cells are sequentially transduced with tdTomato (cell volume marker) and the SORE6 stemness reporter and labeled with red tracker dye. Flow sorting isolates cancer non-stem cells (CnSCs) and cancer stem cells (CSCs) based on SORE6 reporter GFP expression. Cells are then mixed together in a 1:1 ratio. **C)** A lung window is surgically implanted in mice before retro-orbital injection of the 1:1 mixture of CnSCs and CSCs Additionally, mice receive an IV injection of a far-red dextran prior to imaging to label the vasculature. **D)** Representative images showing identification and localization (of tumor cells (yellow arrows) as CnSCs (red) and CSCs (greenish-yellow) relative to blood vessels (white and yellow dashed lines). Cells are either found within the intravascular (upper panels) or the extravascular (lower panels) space. Viability status of cells is determined by size and morphology as survived/died (left panels for CnSCs and CSCs) or died (right panels). **E)** Quantification of the percentage of cancer non-stem cells (CnSCs) versus cancer stem cells (CSC) found extravasated at 32 hours post injection (n=3 mice/group). Unpaired two-tailed Student’s t-test, ***p<0.001. **F)** Quantification of the percentage of CnSCs and CSCs that survive both intravascularly and extravascularly (n=3 mice/group). Unpaired two-tailed Student’s t-test, **p<0.01, ***p<0.001. **G)** Quantification of the percentage of CnSCs and CSCs that both extravasated and survived (n=3 mice/group). Unpaired two-tailed Student’s t-test, ***p<0.001. Panel B and C were created with BioRender.com.

### CSCs and CnSCs do not appreciably interconvert on the hours-to-days frame

Because continuous time-lapse imaging did not reveal any appreciable interconversion between CSCs and CnSCs on the minutes-to-hours time frame, and because this imaging approach inherently restricts the number of cells that can be followed simultaneously, we shifted to the snapshot imaging strategy we previously published (Borriello, Coste et al. 2022). This approach acquires single images every 8 hours and utilizes microcartography (Dunphy, Entenberg et al. 2009) to precisely relocate individual tumor cells for longitudinal tracking over hours-to-days. We also observed minimal interconversion (2 of 172 cells = 1%) during this extended time period. We observed an appreciable number of intra- and extravascular cells at the 32-hour post-injection timepoint which would allow us to look for relative differences in extravasation behavior between CSCs and CnSCs.

Given the common start time of the experiment (i.e. cancer cell injection), and our finding that we could reliably distinguish intravascular from extravascular cells at this stage (**Fig. 2D** and **Supplementary Fig. 6**) without requiring microcartography, we decided to simplify our approach to a single imaging session. This eliminated the need for establishing fiducial coordinate marks and recording xy positions of individual cells, which had limited the total population we could examine. Focusing on this single timepoint (32 hours), enabled rapid and efficient collection of large datasets on the extravasation and survival of CSCs and CnSCs.

### CSCs display an intrinsic advantage in lung extravasation and survival compared to CnSCs

Following the strategy outlined in **Fig. 2D** and **Supplementary Fig.6**, we found that over 53% of CSCs successfully extravasated, compared to only ∼9% of CnSCs (p<0.0004) (**Fig. 2E**). Furthermore, we determined the localization and survival status of CSCs and CnSCs. Interestingly, at 32 hours, all CSCs survived both intravascularly and after extravasation. In contrast, only ∼46% of intravascular CnSCs and ∼31% of extravasated CnSCs remained viable within the lung at 32 hours (**Fig. 2F**). These data show that percentage of all CSCs that have both extravasated and survived is 53.6%, while this percentage is only 2.8% for CnSCs, representing a 19-fold increase (**Fig. 2G**). These results establish that CSCs exhibit a markedly enhanced capacity for extravasation and survival within the lung microenvironment compared to CnSCs, indicating an intrinsic advantage of CSCs in metastatic seeding.

### MenaINV is essential for efficient CSC extravasation in the lung

To investigate whether MenaINV facilitates extravasation of CSCs, we generated MenaINV knockout MDA-MB-231 cell clones using CRISPR/Cas9 (**Supplementary Fig. 7**). Using the two clones (#16 and #19) which produced robust reductions in MenaINV expression in vitro and in vivo (**Supplementary Fig. 8**), we tested their metastatic ability using a spontaneous metastasis model. As expected from the known role of MenaINV in tumor cell intravasation (Pignatelli, Bravo-Cordero et al. 2016) and extravasation (Borriello, Coste et al. 2022), mice bearing tumors derived from MenaINV knock out cells exhibited markedly reduced numbers of lung micrometastases relative to controls (**Supplementary Fig. 9**), indicating that MenaINV deletion impairs the metastatic colonization capacity of these cells. As Clone #16 provided complete abrogation, while Clone #19 still showed some metastatic capacity, we continued with Clone #16, hereafter referred to as MenaINV-KO.

To ensure that the reduced metastatic ability of MenaINV-KO cells was due to the absence of MenaINV and not to the selection of a particular non-metastatic clone, we reintroduced expression of MenaINV in the clone #16 cells by overexpression (hereafter referred to as MenaINV-OE in KO). Quantification confirmed reintroduction at approximately 18-fold above wildtype levels for mRNA and 8-fold for protein **(Supplementary Fig. 10C & 10D)**. We tested the metastatic potential of these cells and found that MenaINV overexpression effectively rescues this capacity (**Supplementary Fig. 11**), confirming the role of MenaINV in metastasis formation.

To evaluate the role of MenaINV in the extravasation of CSCs, we sorted MenaINV wildtype (WT), MenaINV-KO, and MenaINV-OE in KO cells into CSCs and CnSCs and again performed intravital imaging at 32 hours post injection. As before (**Fig. 2E**), the extravasation rate of CnSCs expressing MenaINV WT was low (9%) and deletion of MenaINV did not further decrease this level significantly (7%). However, overexpression of MenaINV in MenaINV-KO in CnSC cells did result in a significant increase in extravasation (31%) (**Fig. 3A**). Meanwhile, for CSCs, deletion of MenaINV led to a significant reduction in the extravasation of CSCs (14%) compared to the WT condition (54%). Overexpression of MenaINV resulted in a jump in extravasation to 73% (**Fig. 3B**).

**Figure 3.**
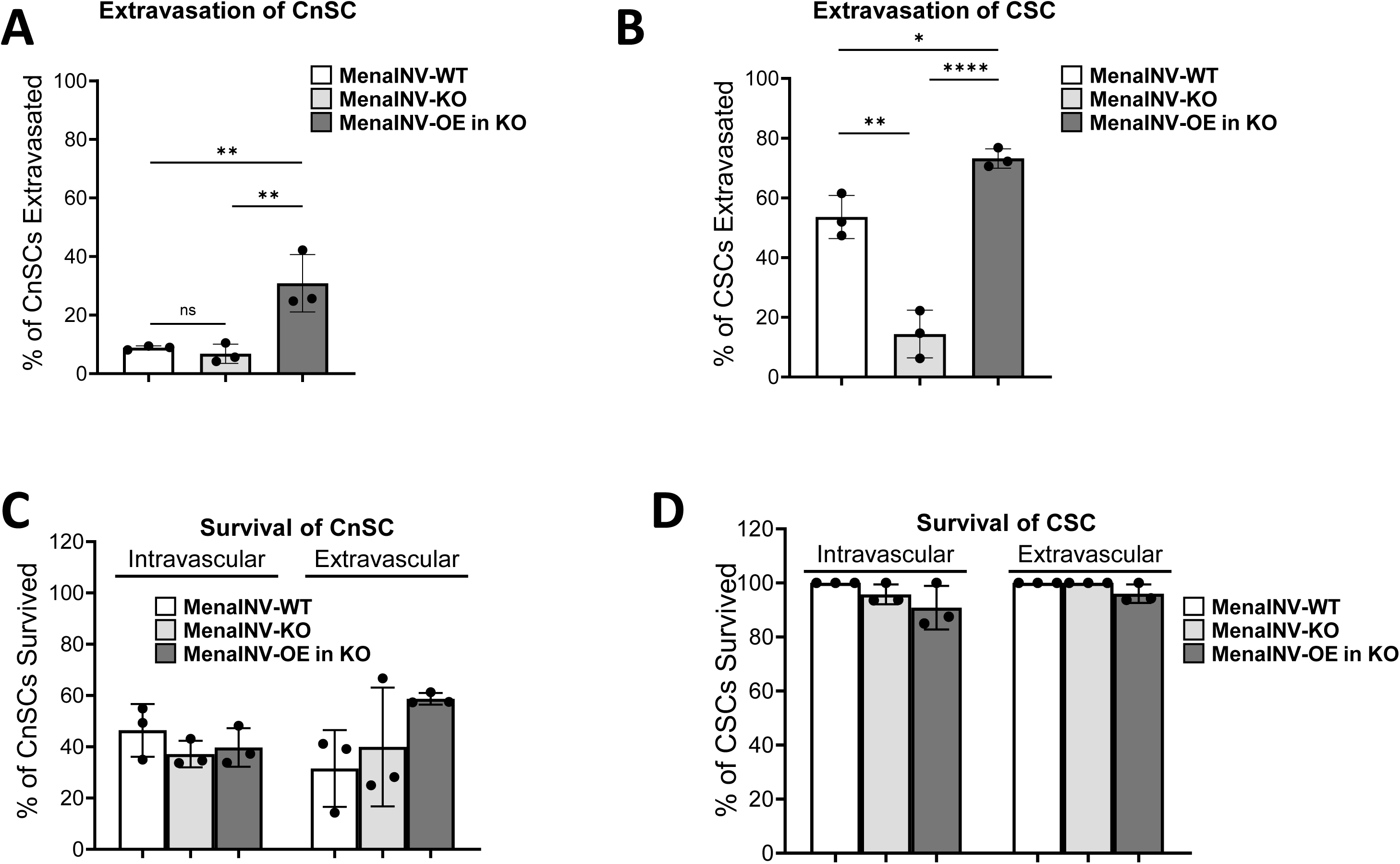
MenaINV is essential for efficient CSC extravasation in the lung, as assessed by intravital lung imaging. **A)** Quantification of the percentage of cancer non-stem cells (CnSCs) found extravasated at 32 hours post injection for MenaINV wild-type (MenaINV-WT), MenaINV knock out (MenaINV-KO), or MenaINV-KO cells that overexpress MenaINV (MenaINV-OE in KO) (n=3 mice/group). One-way ANOVA with Tukey multiple comparison test. **p<0.01. **B)** Quantification of the percentage of cancer stem cells (CSCs) found extravasated at 32 hours post injection for the same groups of cells as in A). (n=3 mice/group). One-way ANOVA with Tukey multiple comparison test, *p<0.5, **p<0.01, ****p<0.0001. **C)** Quantification of the percentage of CnSCs that survive both intravascularly and extravascularly (n=3 mice/group). One-way ANOVA with Tukey multiple comparison test. **D)** Quantification of the percentage of CSCs that survive both intravascularly and extravascularly (n=3 mice/group). One-way ANOVA with Tukey multiple comparison test.

Again, as we observed earlier (**Fig. 2F**), CnSCs with WT MenaINV were impaired in their survival, with only 46% of intravascular and 32% of extravascular cells surviving. Deletion of MenaINV did not substantially affect these levels, resulting in approximately 37% of intravascular cells and 40% of extravasated MenaINV-KO cells remaining viable (**Fig. 3C**). Consistently, we observed that all MenaINV-WT CSCs (100%) survived in the lung, and neither depletion nor overexpression of MenaINV significantly altered this result for cells found in the intravascular (KO=96%, OE=91%) or extravascular (KO=100%, OE=96%) space (**Fig. 3D**).

Taken together, these data demonstrate that MenaINV plays a critical role in enabling CSC extravasation in the lung microenvironment, without impacting their survival.

### MenaINV and stemness cooperate to promote lung metastasis formation

Having determined the role of MenaINV in extravasation and early survival of CnSCs and CSCs, we asked whether the altered MenaINV expression would affect CnSCs and CSCs differently and prevent long term metastatic formation. To evaluate this, we isolated CnSCs or CSCs from MenaINV-WT, MenaINV-KO, and MenaINV-OE in KO cell lines, injected them retro-orbitally (experimental metastases model), and harvested the lungs after the formation of lung metastases.

As expected, mice injected with CSCs expressing MenaINV-WT started to become moribund (weight loss of >15%) after 5 weeks post cell injection, but the mice injected with CnSCs expressing MenaINV-WT, CnSCs with MenaINV-KO, and CSCs with MenaINV-KO did not (**Fig. 4A**). We sacrificed all mice at the 5-week time point and stained their lungs by H&E. We observed no metastases for either group lacking MenaINV, and for the MenaINV-WT CnSCs, as expected (**Fig. 4B**). Also as expected, metastases were observed for the CSCs with MenaINV-WT and MenaINV-OE in KO groups. However, interestingly, we also observed metastasis formation for CnSCs with MenaINV-OE in KO. Quantification and statistical comparison of metastatic area and the number of metastatic foci showed significantly higher levels in MenaINV-WT CSCs compared to MenaINV-WT CnSCs as well as in MenaINV overexpressing cells compared to MenaINV knock out cells, regardless of stemness status at the time of cell injection (**Fig. 4C & 4D**). We previously reported that CnSCs can acquire stemness through interactions with macrophages in tumor microenvironment via Notch signaling in primary breast tumors (Sharma, Tang et al. 2021). This raised the question of whether extravasated CnSCs might similarly acquire stemness in the lung microenvironment, which could explain the unexpected metastatic capacity of MenaINV-overexpressing CnSCs. To test this hypothesis, we performed FLAG immunostaining to detect SORE6 reporter activity in lung metastases at 5 weeks post-injection. Strikingly, we found that 100% of metastatic lesions had acquired a stem-like phenotype after extravasation **(Fig. 4E**). These lesions were indistinguishable from those formed by CSCs (either in MenaINV-WT or MenaINV-OE in KO cells), where stemness marker positivity was also detected in 100% of metastases. These results demonstrate that metastatic lesions formed by MenaINV-overexpressing CnSCs acquired stemness, suggesting that both MenaINV-mediated extravasation and stemness are required for successful metastasis formation.

**Figure 4.**
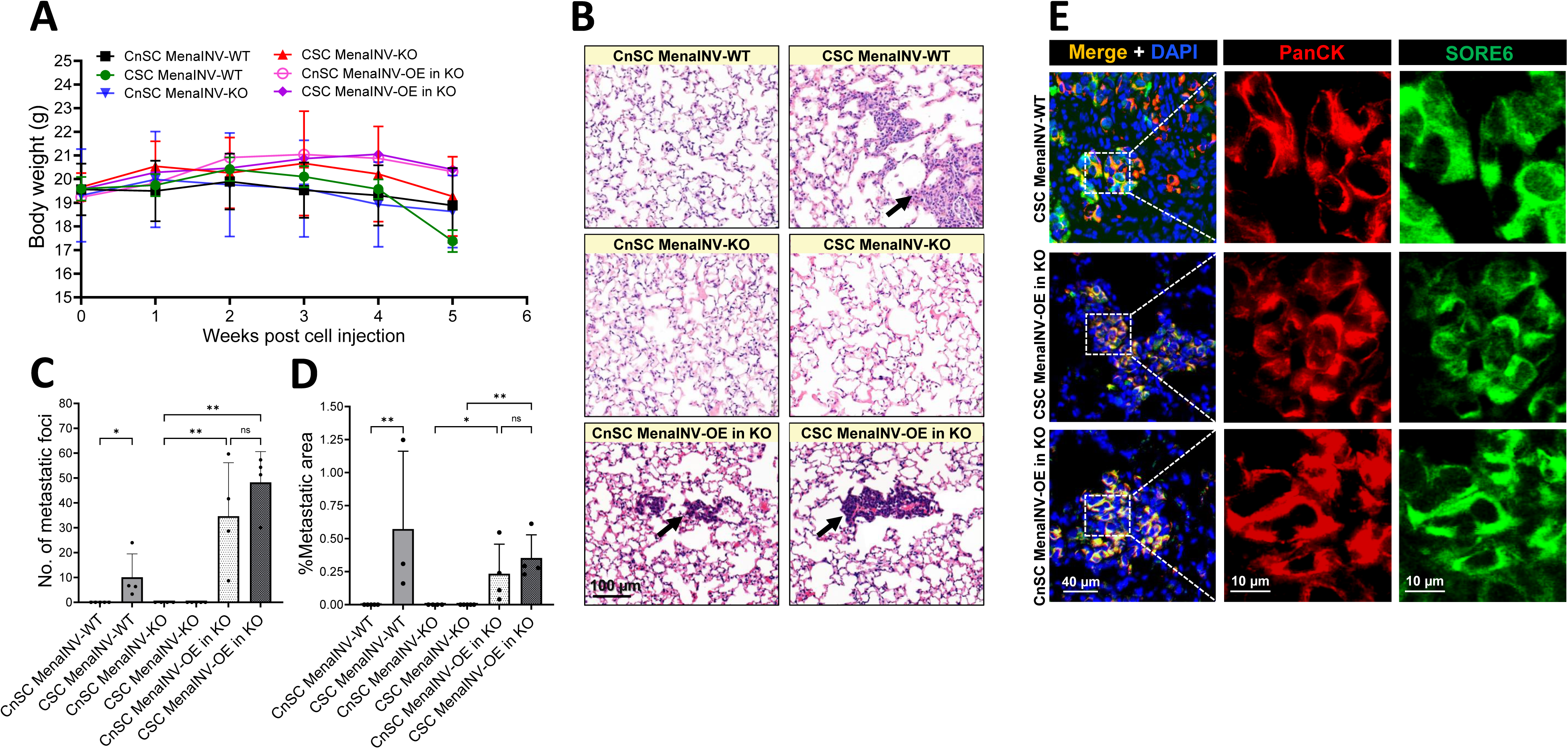
MenaINV and stemness promote lung metastasis formation in a spontaneous metastasis model. **A)** Body weight tracking of mice injected with CnSCs and CSCs expressing MenaINV WT, MenaINV KO, or MenaINV OE in MenaINV KO cells (n=3-5 mice/group). **B)** Representative lung images to show metastases (black arrow) from each experimental group. **C)** Quantification of metastatic area in the lung, assessed by H&E staining (3 sections, 50 µm apart; n=3-5 mice/group). Kruskal-Wallis with Dunn’s multiple comparison test, *p<0.05, **p<0.01. **D)** Quantification of metastatic foci in same sections as in C. (3 sections, 50 µm apart, n=3-5 mice/group). Kruskal-Wallis with Dunn’s multiple comparison test, *p<0.05, **p<0.01. **E)** Representative lung images showing presence of stem cells in the metastatic lesion. pan-CK (red), SORE6 reporter detected via FLAG (green), and nuclei (DAPI, blue) immunofluorescence in CSC MenaINV-WT, CSC MenaINV-OE in KO, and CnSC MenaINV-OE in KO cells. The merged images (PanCK+, FLAG+, DAPI) are shown in the first column. Single-channel images for PanCK and FLAG are shown in the second and third columns, respectively (n=3-4 mice/group). All of the metastatic lesions (100%) from these groups show the presence of CSCs.

## Discussion

Metastasis remains the primary cause of mortality in breast cancer patients (Dillekas, Rogers et al. 2019). While cancer stem cells are well established as drivers of primary tumor formation, their specific role in metastatic formation remains incompletely understood (Amey and Karnoub 2017, Czarnogorski, Czernicka et al. 2025). Metastasis requires tumor cells to complete the sequential steps of the metastatic cascade, including intravasation, extravasation and survival at secondary sites, and finally growth in these locations (Hanahan and Weinberg 2011). Cancer stem cells possess enhanced tumor initiation capacity, increased epithelial to mesenchymal (EMT) status, chemoresistance, and superior survival capabilities compared to the bulk tumor population (Lee, Kim et al. 2025). Our prior work demonstrated CSC progressive enrichment during metastatic progression, with CSCs constituting a surprising 66% of circulating tumor cells, 77% of single cells found both intravascularly and extravascularly in the lung, and 84% of early metastatic lesions (Sharma, Tang et al. 2021). This raised a fundamental question: whether CSC enrichment in early metastases reflects intrinsic functional advantages in specific metastatic steps (such as extravasation and survival) or simply results from their increased representation in circulation due to enhanced intravasation in primary tumors.

Our current study directly addresses this knowledge gap through direct comparison using intravital imaging of cells expressing our validated SORE6 stemness reporter in the live lung. We demonstrate that CSCs possess intrinsic functional superiority over cancer non-stem cells (CnSCs) in extravasation and survival. Most importantly, our mechanistic dissection reveals a hierarchical regulatory framework where stemness acts as a master regulator controlling two functionally distinct pathways: one governing extravasation, and another controlling survival (**Fig. 5**). This framework could provide a molecular explanation for CSC dominance at early stages of dissemination.

**Figure 5.**
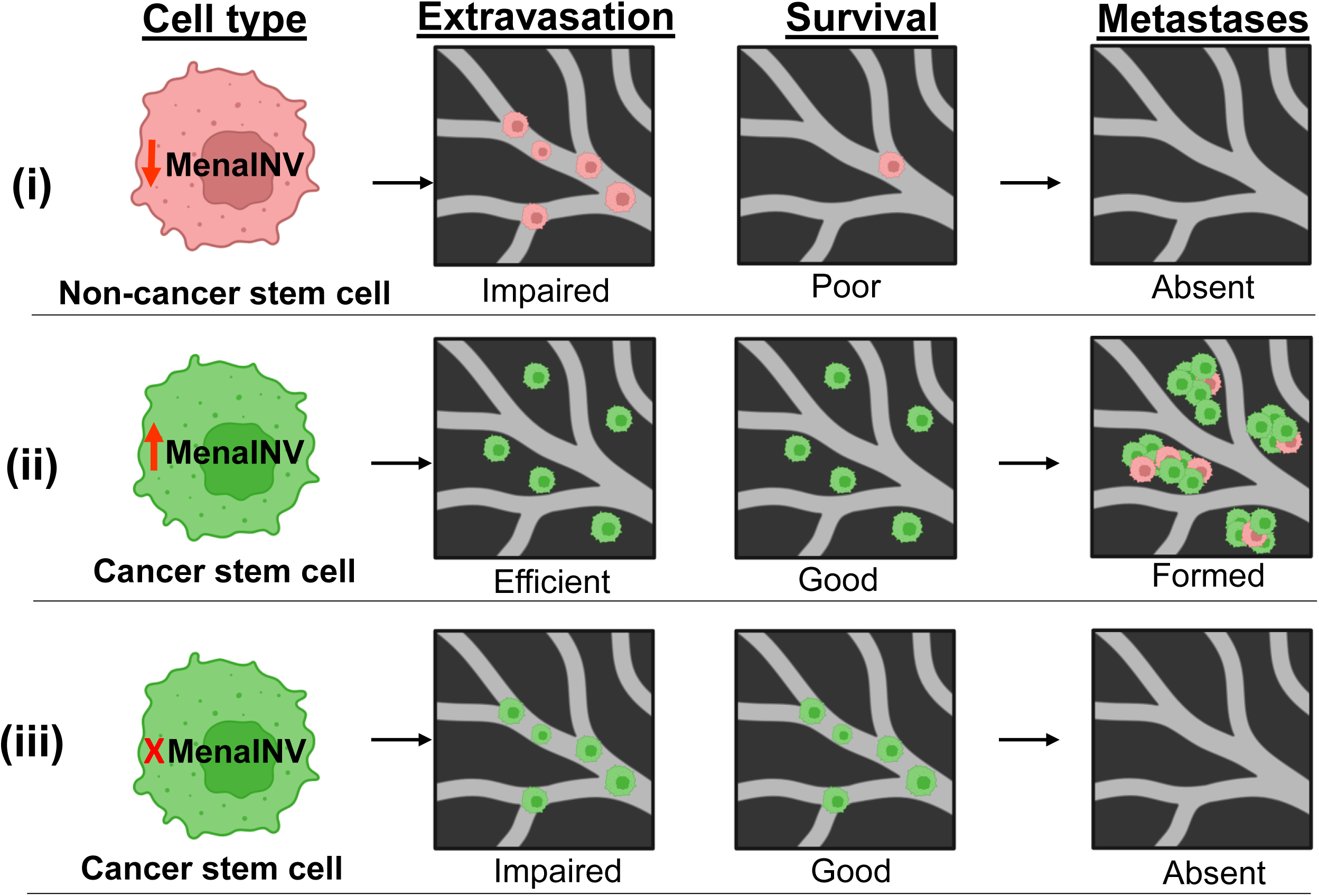
Stemness and MenaINV enable efficient metastatic colonization. Schematic model illustrating the mechanistic relationship between stemness and MenaINV. **(i)** CnSCs express low levels of MenaINV and are impaired in the extravasation ability and do not survive well, resulting in an inability to form metastases. **(ii)** CSCs express high levels of MenaINV and they are able to extravasate efficiently and survive in the lung environment, leading to the formation of metastases. **(iii)** CSCs with MenaINV deletion can survive but lost their ability to extravasate and to form metastases. The image was created with BioRender.com.

The molecular basis for CSC extravasation advantage centers on MenaINV, a splice variant of an actin regulatory protein Mena (Gertler and Condeelis 2011). MenaINV controls multiple processes critical for metastasis, including EMT (Roussos, Balsamo et al. 2011, Roussos, Goswami et al. 2011), chemotaxis (Hughes, Oudin et al. 2015), invadopodia formation (Weidmann, Surve et al. 2016), and transendothelial migration (Pignatelli, Bravo-Cordero et al. 2016). Our findings establish MenaINV as the key downstream effector through which stemness confers extravasation advantages, while survival pathways operate independently of MenaINV.

An important observation from our reintroduction experiments is that MenaINV overexpression (18-fold mRNA and 8-fold protein above endogenous levels, **Supplementary Fig. 10C & 10D**) drives CSC extravasation rates (73%) beyond those observed in wildtype CSCs (54%) (**Fig. 3B**). This dose-dependent enhancement demonstrates that MenaINV functions as a rate-limiting factor for transendothelial migration, with supraphysiological expression conferring proportionally increased extravasation capacity. Critically, despite elevated extravasation, MenaINV overexpression did not enhance survival rates in either CSCs or CnSCs (**Fig. 3C & 3D**), confirming the functional independence of extravasation and survival pathways. These findings establish that while MenaINV levels directly modulate extravasation efficiency, stemness-associated survival advantages operate through entirely distinct molecular mechanisms.

The regulatory hierarchy linking stemness to MenaINV likely involves Sox2 and Sox9, master transcription factors that regulate stemness in both normal development and cancer (Domenici, Aurrekoetxea-Rodriguez et al. 2019). Sox2, whose transcriptional activity is detected by our SORE6 reporter (Tang, Raviv et al. 2015), acts upstream to directly regulate Sox9 expression, establishing a Sox2-Sox9 signaling axis maintaining breast cancer stem cell properties (Domenici, Aurrekoetxea-Rodriguez et al. 2019). This relationship becomes functionally significant through Sox9’s regulatory control over the NF-κB transcriptional network, including Rel, Rela, and NF-κB1 (Christin 2020). Completing this cascade, we recently demonstrated that expression of MenaINV is directly regulated by macrophages through NF-κB signaling in cooperation with Notch pathway activation (Duran, Karagiannis et al. 2023). This suggests a molecular pathway to be explored in future studies (Sox2 → Sox9 → NF-κB → MenaINV) through which stemness facilitates extravasation while maintaining the independence of survival pathway. This hierarchy is in accord with recent findings by us and others that macrophages promote EMT and cancer stemness (Sharma, Tang et al. 2021, Chen, Yang et al. 2022).

Our results expand upon earlier work showing that CSCs possess enhanced metastatic capabilities, including superior invasive properties through enhanced epithelial to mesenchymal transition (EMT) programs (Lei, Teng et al. 2024), enhanced survival mechanisms in metastatic sites (Liu, Taftaf et al. 2019), and enhanced metastatic colonization capacity (Batlle and Clevers 2017, Sharma, Tang et al. 2021). The mechanistic basis for these enhanced invasive properties was established by foundational work demonstrating that the EMT and stem programs are interlinked, with EMT programs generating cells with stem cell properties (Mani, Guo et al. 2008), and CSCs possessing the capacity for invasion and metastasis (Li and Li 2014). However, while these studies established the metastatic advantages of CSCs and the mechanistic link between invasion and stemness, they did not investigate whether these advantages operate through separable molecular pathways or examine the functional independence of invasion versus survival mechanisms.

Our findings reveal, for the first time, the molecular basis of this pathway separation: extravasation operates through a MenaINV-dependent extravasation program, while survival mechanisms function independently of MenaINV. This functional separation has important mechanistic implications. CSCs lacking MenaINV retain their superior survival capacity (96% intravascular survival vs 37% for CnSCs), confirming that survival advantages operate independently of extravasation machinery. Conversely, MenaINV deletion reduces CSC extravasation to levels similar to CnSC (54% vs 14%) (**Fig. 3B**). Furthermore, MenaINV overexpression can rescue extravasation capacity even in CnSCs (31% vs 7% for control CnSCs) (**Fig. 3A**). Most importantly, our long-term colonization studies demonstrate that both stemness and MenaINV are required for successful metastasis formation. Only CSCs expressing MenaINV formed lung metastases at 5 weeks, while CSCs lacking MenaINV and CnSCs (wild type or with MenaINV knockout) failed to establish colonies (**Fig. 4C & 4D**). Remarkably, CnSCs overexpressing MenaINV also formed metastases after acquiring stemness in the lung microenvironment (**Fig. 4E**), confirming that MenaINV enables extravasation but stemness acquisition is essential for colonization.

This study resolves a fundamental question in cancer biology by demonstrating that CSC metastatic advantages reflect intrinsic functional superiority rather than simply increased representation in circulation: a paradigm shift from correlative to mechanistic understanding. By revealing that stemness coordinates metastatic success through functionally separable pathways, our work provides the first mechanistic framework for understanding how CSCs achieve their documented dominance throughout the metastatic cascade. This mechanistic dissection has critical clinical implications for developing treatment approaches for patients with locally advanced and disseminated cancers. Since metastasis remains the primary cause of cancer mortality, targeted prevention of CSC dissemination during pre-surgical systemic treatments and their outgrowth to clinical detectable metastatic foci could provide crucial therapeutic insurance against disease spread. By identifying that both extravasation and survival pathways must be simultaneously disrupted, our findings provide a rational mechanistic basis for developing combination therapies that could halt further spread of metastases and transform metastatic stage from deadly to chronic disease.

## Limitations

While our intravital imaging approach provides direct observation of CSC and CnSC behavior in the lung microenvironment, several experimental limitations should be acknowledged. Our study relies on xenograft models and genetically modified reporter lines, which may not fully recapitulate the complexity of endogenous tumor heterogeneity or the complete human metastatic microenvironment. Additionally, our findings are derived from breast cancer models with a specific focus on lung metastasis, which limits generalizability across other cancer types and metastatic niches. The SORE6 reporter, while validated for detecting Sox2/Oct4 transcriptional activity, represents a surrogate measure of stemness that may not capture the full spectrum of CSC properties or account for context-dependent variations in stemness programs. Future studies should validate these findings across multiple cancer types, metastatic sites, and patient-derived models to establish broader applicability.

While our study establishes that CSCs possess superior survival advantages compared to CnSCs in the lung microenvironment, the molecular mechanisms underlying this enhanced survival remain largely unknown. Our analysis focused on short-term survival dynamics (within hours to days), demonstrating MenaINV-independent survival pathways that operate distinct from extravasation mechanisms. However, we previously showed that stemness, as identified via CD44 expression, conveys a long-term survival advantage to disseminated tumor cells in the lung over extended periods (Liu, Taftaf et al. 2019). It remains unclear whether the same molecular mechanisms and signaling pathways that regulate short-term CSC survival also govern long-term persistence and dormancy of disseminated tumor cells in the pulmonary microenvironment. The temporal dynamics of survival mechanisms may involve distinct molecular programs, with early survival potentially driven by different pathways than those required for long-term persistence and eventual outgrowth. Further investigation is needed to elucidate the molecular underpinnings governing both short-term and long-term survival of CSCs in the lung, including the potential transitions between survival states and the microenvironmental factors that influence these temporal survival programs.

## Conclusion

This work establishes a direct mechanistic link between the ability of cancer stem cells to enhance MenaINV-mediated extravasation while providing survival advantages for cancer cells resulting in efficient metastatic colonization in the lung. By clarifying the functional interplay between these pathways, it advances the prospects for rational intervention strategies targeting both cancer stemness and MenaINV to limit breast cancer metastasis.

## Methods

### Cell culture

The MDA-MB-231 human breast cancer cell line was obtained from the American Type Culture Collection (ATCC). Cell line identity was validated by short tandem repeat (STR) profiling (Laragen Corp.) following initial expansion and passaging. Both parental and genetically manipulated MDA-MB-231 cells were cultured in Dulbecco’s Modified Eagle Medium (DMEM) supplemented with 10% fetal bovine serum (FBS) and 1x penicillin-streptomycin under standard conditions (37°C, 5% CO_2_).

### Introducing tdTomato as a cell volume marker

Parental MDA-MB-231 cells, along with all manipulated cells, were transduced with a lentiviral vector encoding tdTomato (Addgene plasmid #112579) to enable fluorescent labeling and subsequent tracking of cell volume. tdTomato-expressing cells were sorted through fluorescence-activated cell sorting (FACS) consecutively to ensure a uniformly labeled population.

### CRISPR/Cas9 mediated knockout of MenaINV

The guide RNA (gRNA) – GAGCAAGGTTACTGCTACCC, targeting the MenaINV was designed using the Integrated DNA Technologies (IDT) online CRISPR-Cas9 gRNA design tool. MDA-MB-231-tdTomato cells were transfected using the IDT ribonucleoprotein (RNP) lipofection method with either a non-targeting control gRNA (NT gRNA) or MenaINV gRNA, tracrRNA conjugated to ATTO647 dye, and recombinant Cas9 protein. Transfection was carried out for 48 hours. At 24-hours post-transfection, cells were sorted by flow cytometry to isolate tdTomato+ATTO647+ (double-positive) cells. Single-cell clones were established by seeding one tdTomato+ATTO647+ cell per well in 96-well plates. After 8–10 days of expansion, individual colonies were screened for genome editing efficiency using a T7 endonuclease I (T7EI) and polymerase chain reaction (PCR) assay. Two of the candidate knockout clones, designated as #16 and #19, were further validated by Sanger sequencing and ICE analysis confirming targeted deletion in MenaINV specific exon. Clone #16 exhibits a clean biallelic knockout with two predominant indels, reflecting a monoclonal population with predictable gene disruption, as indicated by a high knockout score and strong model fit (R²). In contrast, Clone #19 represents multiple diverse indels, suggesting genetic heterogeneity, likely from polyploidy or mosaicism, hence we picked Clone #16 for subsequent experiments investigating extravasation and metastatic growth. However, we confirmed deletion of MenaINV in both Clones #16 and #19 by quantitative RT-PCR (qRT-PCR) in cells and also by Western blot analysis in primary breast tumor samples.

### Reintroduction of MenaINV in MenaINV KO cells

The pHTN-HaloTag-MenaINV construct was created by subcloning eGFP-MenaINV from the pMSCV-eGFP-MenaINV plasmid (Philippar, Roussos et al. 2008) into the pHTN-HaloTag CMV-neo vector using Pvul and NotI insert sites (Pignatelli, Bravo-Cordero et al. 2016). MenaINV KO cells were transfected with a pHTN-HaloTag-MenaINV plasmid using Lipofectamine 3000 and P3000 reagent in low serum (5%) culture medium and incubated for 24 hours in incubator supplied with 5% CO_2_. After 24 hours of incubation, culture media was replaced with 10% serum medium. Stable cells were selected on 1 mg/mL G418 antibiotic. Expression of MenaINV was verified by qRT-PCR and Western blot analysis in cell culture.

### Stemness reporter

To identify cancer stem cells (CSCs), we employed the second-generation SORE6 fluorescent stemness reporter, as previously described (Sharma, Tang et al. 2021). The SORE6 reporter consists of six tandem repeats of a composite Sox2/Oct4 response element upstream of a minimal cytomegalovirus (minCMV) promoter, which drives the expression of a destabilized CopGFP (dsCopGFP) fluorescent reporter (Tang, Raviv et al. 2015) (**Fig. 2A and Supplementary Fig. 1A**). As the native dsCopGFP fluorescence signal is lost during fixation, this construct includes an N-terminal 3xFLAG epitope tag fused to dsCopGFP, enabling reliable detection by immunofluorescence in formalin-fixed paraffin-embedded (FFPE) tissues. In addition, a C-terminal truncated CD19 surface marker is also incorporated into the reporter construct, allowing for FACS separation of successfully transduced cells. To establish a threshold for GFP expression and eliminate background fluorescence, a control construct (minCMV control) lacking the SORE6 response elements (**Supplementary Fig. 1B**) was used in parallel for FACS gating and multiphoton intravital imaging. For intravital imaging, control mice injected with minCMV control cells were imaged to establish background GFP signal levels (**Supplementary Fig. 5**).

### Fluorescence-activated cell sorting

Flow cytometry-based fluorescence-activated cell sorting (FACS) was employed across multiple experimental workflows to isolate MDA-MB-231 cells based on specific fluorescent markers. Cells expressing tdTomato were selected to track cell volume, while double-positive tdTomato+ATTO647+ cells were isolated during CRISPR/Cas9-mediated genome editing.

For experiments involving the SORE6 stemness reporter, cells were sorted into SORE6- and SORE6+ populations based on GFP expression. Because the SORE6 construct includes a truncated CD19 surface marker, transduced cells were first gated as CD19+ using an APC-conjugated anti-CD19 antibody to ensure selection of reporter-positive cells. Details on antibodies utilized are provided in **Supplementary Table 1.**

The threshold for GFP signal above which cells are identified as CSCs was established using a minCMV control construct (called tdTomato+CD19+minCMV control) which lacks Sox2/Oct4 response elements and therefore only generates background fluorescence. Based on this threshold, tdTomato+CD19+SORE6-cells (GFP fluorescence below threshold) were classified as CnSCs, and tdTomato+CD19+SORE6+ (GFP fluorescence above threshold) as CSCs. To generate pure populations of CSCs and CnSCs, we first sorted CD19+ cells to ensure successful transduction with the SORE6 reporter construct, and then sorted the top ∼10% (SORE6+, CSCs) and lowest 10% (SORE6-, CnSCs) GFP expressing cells from this CD19+ population.

### Antibodies and primers

Chicken anti-MenaINV antibodies were generated by Covance, as previously described (Pignatelli, Bravo-Cordero et al. 2016). A complete list of primary and secondary antibodies used in this study is provided in **Supplementary Table 1**. The primers for qRT-PCR were obtained from IDT and their sequences are listed in **Supplementary Table 2**.

### qRT-PCR

Total RNA was extracted from cultured cancer cells using the RNeasy Mini Plus Kit (Qiagen, Cat# 74134) according to the manufacturer’s protocol. RNA was eluted in RNase-free water and quantified using a NanoDrop spectrophotometer (Thermo Fisher Scientific). First-strand complementary DNA (cDNA) synthesis was performed using the iScript cDNA Synthesis Kit (Bio-Rad, Cat# 1708891), following the manufacturer’s instructions. Quantitative real-time PCR (qRT-PCR) reactions were carried out using the Power SYBR Green Master Mix (Thermo Fisher Scientific, Cat# 4367659) on the QuantStudio 3 Real-Time PCR System (Applied Biosystems). Reactions were performed in triplicate using a 96-well plate and run for 40 amplification cycles. Gene expression levels were normalized to GAPDH as the internal control and were calculated as relative expression using the log2ddCT method.

### In vitro IF staining of CnSCs (SORE6-) and CSCs (SORE6+) with anti-MenaINV

MDA-MB-231 cells transduced with the SORE6 reporter were sorted into SORE6- and SORE6+ cell populations and cultured on Millicell EZ slides (Millipore Cat# PEZGS0816), fixed with 3.7% paraformaldehyde in DPBS for 15 min at room temperature. After washing 3x for 5 min each with DPBS, the fixed cells were permeabilized with 0.1% Triton X-100 in DPBS for 10 min followed by blocking with 2.5% BSA + 10% normal goat serum in DPBS for 1 h at room temperature. The slides were incubated with MenaINV antibody at 4°C overnight. Slides were washed 3x for 5 min with DPBS and incubated with corresponding secondary antibodies (**Supplementary Table 1**). Slides were imaged on 3D Histech Pannoramic 250 Flash III digital whole slide scanner, using a 20x 0.8 NA air objective lens and were analyzed for mean intensity of MenaINV and SORE6 reporter (via GFP and FLAG) using the Visiopharm image analysis software, Vis (Visiopharm, Hørsholm, Denmark).

### In vivo determination MenaINV in primary breast tumor by IF staining

MDA-MB-231-tdTomato+SORE6 cells were injected into 4^th^ mammary fat pad of NOD-SCID mice to develop primary breast tumors. Primary tumors were harvested at ∼60 mm^3^ of size and FFPE blocks were made. 5 µm tissue sections were cut for staining with anti-MenaINV and anti-FLAG tag to detect SORE6 reporter. The unstained slides were first baked at 55°C for 1 hour followed by deparaffinization in Xylene and rehydration in a graded series of ethanol. Antigen unmasking was performed in antigen retrieval pH 9 buffer (Vector Labs, Cat# H-3301) in a conventional steamer. The sections were washed 3x for 5 min each in PBST and incubated in blocking buffer (1% BSA + 10% normal goat serum in 1x PBST) at room temperature for 1 h. The sections were incubated with antibody cocktails containing anti-MenaINV (0.25µg/mL, AE1071, AP-6) and anti-FLAG (1:200, Biolegend, Cat# 637301) overnight at 4°C. After washing with 3x for 5 min each in PBST, another cocktail of corresponding secondary antibodies **(Supplementary Table 1)** was added onto the sections for one hour at room temperature. The IF stained slides were images on the 3D Histech Pannoramic 250 Flash III digital whole slide scanner using a 20x 0.8 NA air objective lens and digital whole slides were analyzed for relative expression of MenaINV in SORE6-FLAG expressing cells using the Visiopharm image analysis software.

### Quantification of MenaINV, SORE6-GFP and SORE6-FLAG

Regions of interest (ROIs) on digital whole slides were manually defined to include all cells (in vitro) or tumor areas that met pathological quality criteria, including tumor adequacy and absence of necrosis, inflammation, tissue folds, or retraction artifacts (in vivo). For each digital whole slide, the total ROI area was chosen to span and area of at least ten high-power (20x) fields of view (600×880 μm²). Image analysis was performed by generating a nuclear mask from the DAPI channel. A cytoplasmic mask was then created by expanding the nuclear mask by 3 pixels and subtracting the nuclear region to isolate the perinuclear cytoplasmic compartment. The perinuclear cytoplasmic mean intensity was quantified for SORE6-FLAG, SORE6-GFP, and MenaINV. Mann–Whitney U tests were applied to compare the mean intensities of MenaINV and SORE6-GFP (in vitro) and Pearson correlation analyses were performed to assess the association between SORE6-FLAG and MenaINV expression (in vivo).

### Identification of cancer stem cells (CSCs) in the fixed lung tissues by IF staining

Lungs collected from experimental metastasis models were co-stained with an antibodies against human specific pan-cytokeratin (PanCK) and FLAG. All subsequent staining steps followed the same protocol as for primary tumor IF staining. Secondary antibodies used are listed in **Supplementary Table 1**. Slides were imaged with a 3D Histech Pannoramic 250 Flash III digital whole slide scanner, using a 20x 0.8 NA air objective lens and analyzed for the presence of CSCs in metastatic growths.

### Western blotting

Primary breast tumor (∼50 mg) tissue or culture cells were homogenized in 100 μL of lysis buffer containing 1% IGEPAL CA-630, 1% Triton X-100, 0.5% sodium deoxycholate (all from Sigma-Aldrich), and Complete™ Protease Inhibitor Cocktail (Cat# 11836170001, Roche, Sigma-Aldrich). Homogenates were incubated on ice for 30 minutes and subsequently centrifuged at 5,000x g for 5 minutes at room temperature. The resulting supernatant was collected and supplemented with 20% glycerol. Protein concentration was determined using the Bradford Protein Assay (Bio-Rad Laboratories). For SDS-PAGE, 50 µg of total protein was mixed with loading dye, separated by electrophoresis, and transferred onto a nitrocellulose (NC) membrane. Membranes were blocked for 1 hour at room temperature in 5% non-fat dry milk prepared in Tris-buffered saline containing 0.05% Tween-20 (TBST). Following blocking, membranes were sectioned between the ∼50 and ∼75 kDa molecular weight markers and incubated overnight at 4°C with one of the following primary antibodies diluted in blocking buffer: chicken anti-MenaINV (0.25µg/mL, AE1071, AP-4) or Mouse anti-Pan Mena (1:1000, EMD Millipore Cat# MAB2635) or Mouse anti-HaloTag (1:1000, Promega, Cat# Cat# G921A) or Rabbit anti-actin (1:2000, Sigma-Aldrich, Cat# A2066). After washing with TBST, membranes were incubated with the appropriate species-specific secondary antibodies (listed in **Supplementary Table 1**) for 1 hour at room temperature in blocking buffer. Protein bands were visualized either by chemiluminescence on X-ray film (Thermo Fisher Scientific) or using the Odyssey Classic Infrared Imaging System (LI-COR Biosciences).

### Identification of disseminated tumor cells in the fixed lung tissues by IF staining

Lungs collected from spontaneous or experimental metastasis models were co-stained with an antibody cocktail against human specific pan cytokeratin and FLAG to detect the SORE6 reporter. All subsequent staining steps followed the same protocol as for primary tumor IF staining. Secondary antibodies used are listed in **Supplementary Table 1**. Slides were imaged with a 3D Histech Pannoramic 250 Flash III digital whole slide scanner, using a 20x 0.8 NA air objective lens and analyzed for disseminated tumor cells or metastatic growth.

### Animal models

All procedures were conducted in accordance with the National Institute of Health regulation concerning the care and use of experimental animals with the approval of Albert Einstein College of Medicine Animal Care and Use Committee (IACUC). Female mice aged 8-10 weeks old were used throughout.

For the spontaneous metastasis model, NOD SCID mice (Jackson Labs, strain #001303) were injected orthotopically in the 4^th^ and 9^th^ mammary fat pads with 2×10⁶ MDA-MB-231 cells resuspended in 50 µL of PBS and collagen (Sigma-Aldrich Cat#C3867-1VL) at a 1:1 ratio. Experiments included **i)** validation of CRISPR/Cas9 mediated MenaINV deletion using SORE6 expressing MenaINV control (gRNA), MenaINV KO, or MenaINV KO with MenaINV overexpression cells, as described in **CRISPR/Cas9 mediated knockout of MenaINV** section, and **ii)** assessment of the correlation between stemness and MenaINV in primary tumors formed from SORE6 reporter cells.

For experimental metastasis models, Rag2 KO mice (Jackson Labs, strain #008449) received retro-orbital injection of 2×10⁶ cells resuspended in 200 µL PBS. Sorted cell populations included SORE6 expressing control (gRNA) and MenaINV knockout or overexpression variants in a 1:1 ratio of CSC to CnSCs.

For extravasation studies, Rag2 KO x MacBlue mice bearing a permanent lung imaging window (Entenberg, Voiculescu et al. 2018, Borriello, Coste et al. 2022) received similar retroorbital injection of sorted cell populations as above for intravital tracking of CnSC and CSC extravasation in the lung, described in further detail below.

### Surgery to implant a Window for High-Resolution Intravital Imaging of the Lung (WHRIL)

Prior to surgery, mice were anesthetized with isoflurane and received pre-emptive analgesia to minimize procedural pain. After induction of anesthesia, a permanent Window for High-Resolution Intravital Imaging of the Lung (WHRIL) was surgically implanted under aseptic conditions, as described in (Entenberg, Voiculescu et al. 2018, Borriello, Traub et al. 2021, Borriello, Coste et al. 2022). Briefly, the skin over the upper chest was sterilized and depilated; mice were intubated with a catheter, ventilated, and positioned in the right lateral decubitus position on a warm surgical platform. A circular incision of ∼10 mm was made through the skin on the upper chest, muscles overlaying the ribs were removed, and a 5 mm opening was created through the rib cage to expose the lung. A passivated window frame was fitted and sutured in place, the lung was inflated, and a coverslip coated with a thin layer of adhesive was placed securely onto the lung tissue. Skin surrounding the incision was also sutured within the window frame to prevent infection and excess air removed from the thoracic cavity using an insulin syringe inserted through the diaphragm. Mice were allowed to recover nearly completely from surgery on 100% oxygen, at which point they were extubated and administered an additional dose of analgesia. Mice were then moved to clean cages and provided with antibiotics in their drinking water.

### Intravital imaging

SORE6 reporter cells were FACS sorted into CnSCs (SORE6-) and CSCs (SORE6+) based on GFP expression, followed by short term culture in separate culture dishes to allow cells to recover. We frequently observed drift in tdTomato expression in the experimental cells. To minimize variability in fluorescence intensity and ensure reliable identification, tumor cells were additionally labeled with a red cell tracker dye (Invitrogen, Cat# C34552) right before the injection. 2.5×10^5^ CnSCs and 2.5×10^5^ CSCs were mixed together and injected retro-orbitally into WHRIL bearing Rag2 KO MacBlue mice followed by intravital imaging of lung, as previously described (Entenberg, Wyckoff et al. 2011, Entenberg, Pastoriza et al. 2017). Mice were kept under continuous anesthesia using 1-2% isoflurane during imaging. The physiological temperature was maintained using a built-in environment heat enclosure, and a pulse oximeter (PhysioSuite, Kent Scientific) was used to monitor the vitals. 300 µg of 150 kDa far-red dextran (Biotium, Cat# 80132) dissolved in 100 µL PBS was administered retro-orbitally to visualize the blood vasculature. All images were captured in 16 bit using a 25x 1.05 NA objective lens.

#### Analysis of intravascular and extravascular disseminated cancer cells

Cancer cells were classified as either intravascular or extravascular (extravasated) based on their spatial localization relative to the vasculature. CnSCs (SORE6-) and CSCs (SORE6+) were distinguished by their fluorescence profile: CnSCs exhibited red fluorescence due to constitutive expression of tdTomato combined with red cell tracker staining, whereas CSCs displayed greenish-yellow fluorescence resulting from the additional expression of the SORE6 reporter. During intravital imaging each mouse injected with tumor cells and visual inspection through the microscope ocular was used to count the total number of cells in a field of view (position). In each experimental mouse, 30-85 fields of view containing a total of 100-700 tumor cells were recorded. These cells were identified as CSCs or CnSCs by their fluorescence signal and then further identified either as identified as intravascular or extravascular, and viable or non-viable.

## Statistical analysis

All analyses were conducted using GraphPad Prism version 10.0.2 (Dotmatics, Boston, MA).

## Acknowledgements

We would like to acknowledge Mary Chen from Translational Pathology Core for providing Mena null cells; the Histology & Comparative Pathology Core Facility at Einstein College of Medicine for their help with tissue processing, sectioning and H&E staining; the Analytical Imaging Facility at Einstein College of Medicine for digital whole slide scanning, The laboratories of Drs. Lalage Wakefield and Binwu Tang’s at the National Cancer Institute for their collaboration in the use and interpretation of the SORE6 reporter construct.

This research was supported by NIH R01 CA255153, P01 CA257885, S10 OD019961, and P30 CA013330; Sir Henry Wellcome Trust 221647/Z/20/Z; the Evelyn Gruss Lipper Charitable Foundation, the Helen & Irving Spatz Family Foundation, the Gruss-Lipper Biophotonics Center, and the Integrated Imaging Program for Cancer Research.

## Author contributions

Conceptualization – D.E., J.S.C., M.N., M.H.O., and Y.J.; Methodology – B.F.K., C.L.D., D.E., J.L., J.S.C., M.H.O., M.N., M.D., N.D.B., P.P., R.E., and Y.J.; Formal analysis – D.E., M.N., S.S., and Y.J.; Software – D.E.; Investigation – D.E., J.S.C., M.H.O., M.N., and Y.J.; Writing – D.E., J.S.C., M.H.O., and M.N.; Funding acquisition – D.E., M.H.O., and J.S.C.; Resources – D.E., J.S.C., and M.H.O.; Supervision – D.E., J.S.C., and M.H.O.

