## Supplementary data for "Metastatic dissemination of breast cancer stem cells requires MenaINV for lung extravasation but not survival"

**Table of contents**

Supplementary Table 1

Supplementary Table 2

Supplementary Figure 1

Supplementary Figure 2

Supplementary Figure 3

Supplementary Figure 4

Supplementary Figure 5

Supplementary Figure 6

Supplementary Figure 7

Supplementary Figure 8

Supplementary Figure 9

Supplementary Figure 10

Supplementary Figure 11

**Supplementary Table 1. List of antibodies used in the study.**

| Target | Primary antibody | Secondary antibody | Technique done |
|---|---|---|---|
| CD19 | Mouse anti-Human CD19 APC, Biolegend Cat# 392504 (dilution 1:20) | Not applicable | Flow sorting |
| MenaINV | Chicken anti-MenaINV AE1071 AP-4 or AP-6 (0.25μg/mL) | Rabbit anti-Chicken HRP, Invitrogen Cat# SA1-9509 (1:5000)<br><br>Donkey-anti-Chicken IRDye 800, Licor Cat# 925-32218 (1:5000)<br><br>Goat-anti-Chicken AF488, Invitrogen Cat# A32931 (1:600)<br><br>Goat-anti-Chicken AF555, Invitrogen Cat# A32932 (1:500) | Western Blotting<br><br>Western Blotting<br><br>IF Staining<br><br>IF Staining |
| pan-Mena | Mouse anti-pan-Mena, EMD Millipore Cat# MAB2635 (1:1000) | Donkey-anti-Mouse IRDye 680, Licor Cat# 926-68072 (1:5000) | Western Blotting |
| pan-Cytokeratin | Mouse anti-Human pan-Cytokeratin, Origene Cat# CF190321 (1:200) | Goat anti-Mouse AF488, Invitrogen Cat#A11001 (1:600) | IF staining |
| FLAG tag (DYKDDDK) | Rat anti-FLAG tag (DYKDDDDK-tagged mouse Langerin), Biolegend, Cat# 637301(1:200) | Goat anti-Rat AF647, Invitrogen Cat# A21247 (1:1000) | IF staining |
| HaloTag | Mouse HaloTag, Promega Cat# G921A (1:1000) | Donkey-anti-Mouse IRDye 680, Licor Cat# 926-68072 (1:5000) | IF staining |
| Actin | Rabbit anti-Actin, Sigma-Aldrich Cat# A2066 (1:2000) | Goat anti-Rabbit HRP, Invitrogen Cat# 65-6120 (1:5000)<br><br>Donkey-anti-Rabbit IRDye 680, Licor Cat# 926-68073 (1:5000) | Western Blotting<br><br>Western Blotting |
| GAPDH | Mouse anti-GAPDH, Abcam Cat# AB8245 (1:5000) | Donkey-anti-Mouse IRDye 680, Licor Cat# 926-68072 (1:5000) | Western Blotting |

**Supplementary Table 2. Sequence of human primers used in the study.**

| Gene | Primer sequences 5'→3' |
|---|---|
| Sox2 | F: GGGGAAAGTAGTTTGCTGCCTCT<br>R: TGCCGCCGCCGATGATTGTT |
| Oct4 | F: CCTGAAGCAGAAGAGGATCACC<br>R: AAAGCGGCAGATGGTCGTTTGG |
| Sox9 | F: GTACCCGCACTTGCACAAC<br>R: TCTCGCTCTCGTTCAGAAGTC |
| CD44 | F: CCAGAAGGAACAGTGGTTTGGC<br>R: ACTGTCCTCTGGGCTTGGTGTT |
| CD133 | F: CACTACCAAGGACAAGGCGTTC<br>R: CAACGCCTCTTTGGTCTCCTTG |
| ALDH1 | F: CGGGAAAAGCAATCTGAAGAGGG<br>R: GATGCGGCTATACAACACTGGC |
| P63 | F: CAGGAAGACAGAGTGTGCTGGT<br>R: AATTGGACGGCGGTTCATCCCT |
| ITGB4 | F: AGGATGACGACGAGAAGCAGCT<br>R: ACCGAGAACTCAGGCTGCTCAA |
| MenaINV | F: AGAGGATGCCAATGTCTTCG<br>R: TACATCGCAAATTAGTGCTGTC |
| Mena11a | F: CAACAAGAAAACCTTGGGAAA<br>R: GGACCTGTTGTCAAAAACAATCT |
| Pan-Mena | F: AGGCTGAAGCAGGACATTTT<br>R: TGCTCAGTTCCTGCCTGAT |
| GAPDH | F: CGACCACTTTGTCAAGCTCA<br>R: CCCTGTTGCTGTAGCCAAAT |

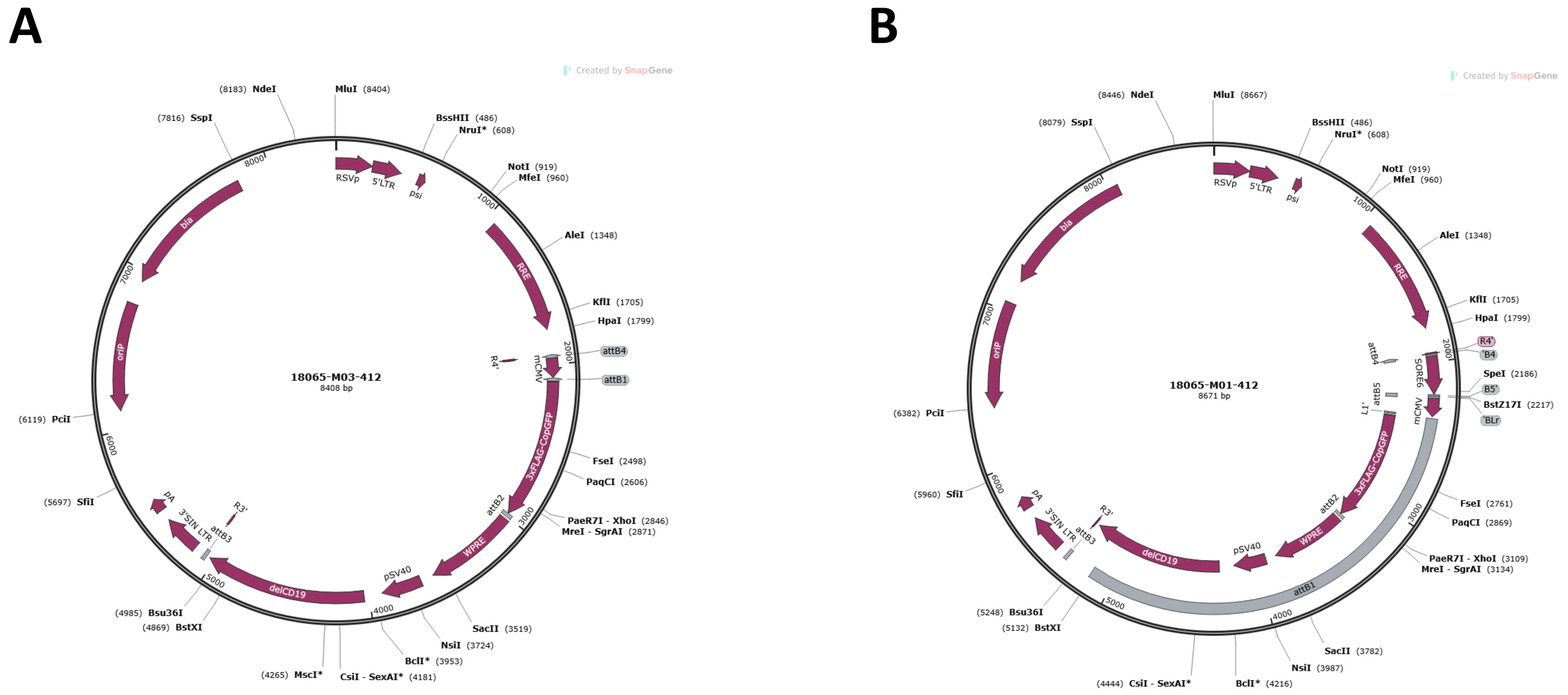


**Supplementary Figure 1. Maps of minimal CMV and SORE6 reporter construct. A)** represents all the components of minimal CMV (minCMV) construct lacking SORE6 response elements. **B)** Represents the full map of SORE6 reporter construct possessing the additional SORE6 (Sox2 and Oct4 response element) compared to minCMV construct.

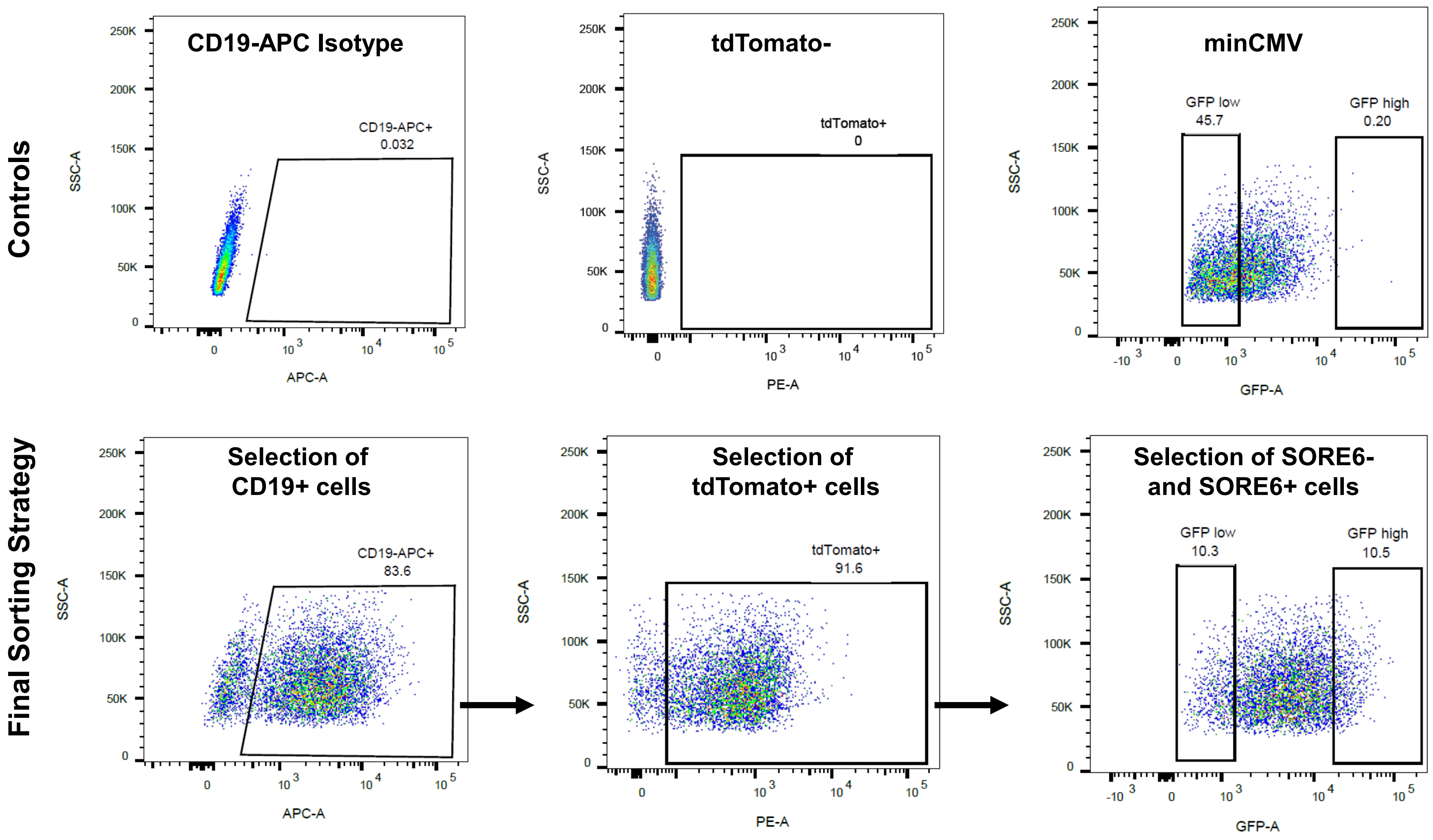


**Supplementary Fig. 2**

**Supplementary Figure 2. Representative gating strategy for flow sorting MDA-MB-231-tdTomato SORE6- and MDA-MB-231-tdTomato SORE6+ cells.** Top Row: Appropriate flow gates were determined by analyzing control cells (Top Row) consisting of CD19 isotype control (APC), tdTomato negative cells, and cells transfected with the minCMV control construct. GFP-high gate was set above all minCMV control cell signal. Final sorting strategy (Bottom Row) collected the top and bottom 10% of SORE6 reporter cells as CSCs and CnSCs. Abbreviations: CSC, cancer stem cell; CnSC, cancer non-stem cell.

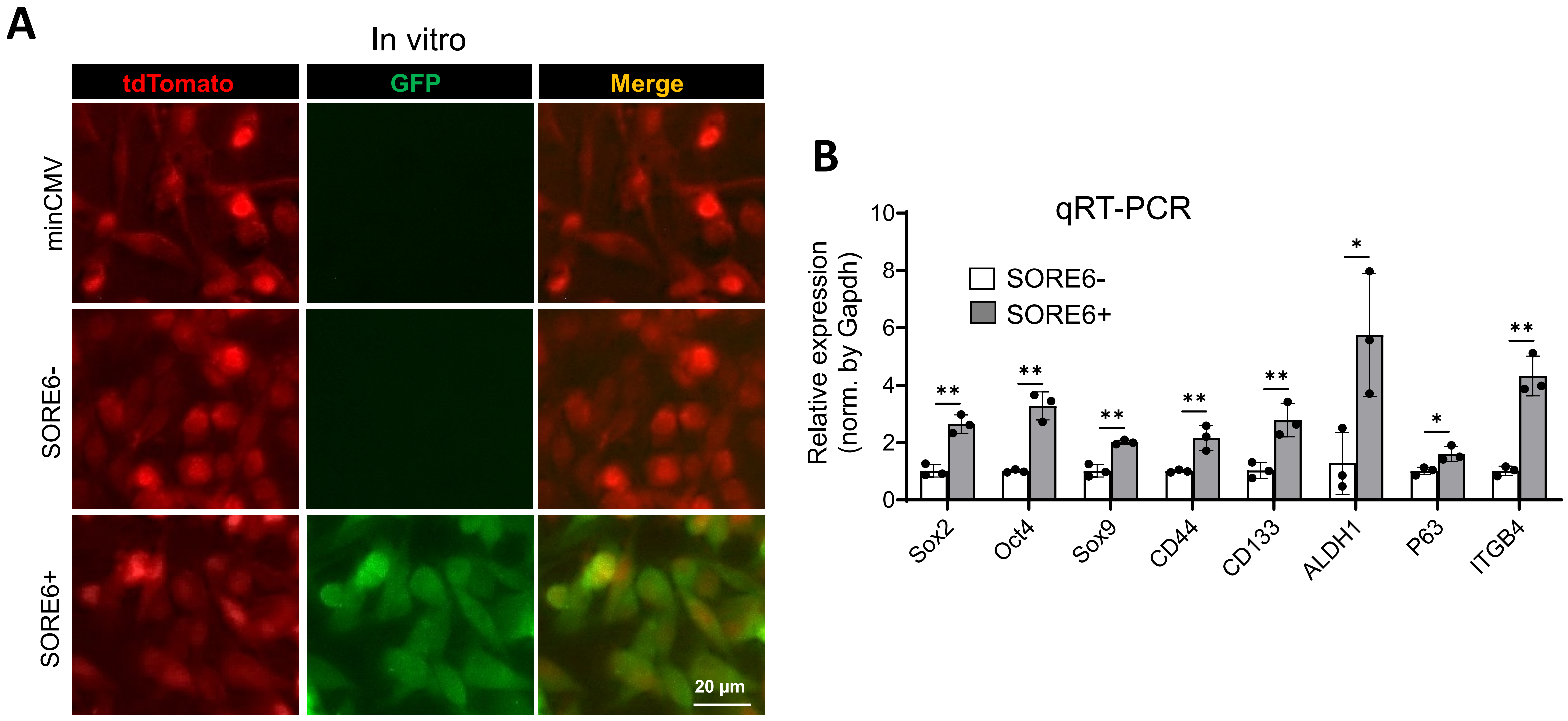


**Supplementary Fig. 3**

**Supplementary Figure 3. Validation of tdTomato and SORE6 reporter expression in sorted cell populations. A)** Representative images showing fluorescence expression of sorted MDA-MB-231 cells. Top Row: minCMV control cells exhibit only tdTomato fluorescence. Middle Row: SORE6- cells also show exclusive tdTomato fluorescence, confirming successful identification of the non-stem population. Bottom Row: SORE6+ cells display both tdTomato and GFP fluorescence, confirming successful identification of the stem population. **B)** Quantitative RT-PCR comparing transcription factors in sorted SORE6- versus SORE6+ cells. SORE6+ cells have elevated expression of Sox2, Oct4, Sox9, CD44, CD133, ALDH1, P63, and ITGB4. Error bars = mean±SEM (n=3 PCR assays), normalized to GAPDH and quantified using the log2ddCT method. Unpaired two-tailed Student's t-test, *$p<0.05$, **$p<0.01$.

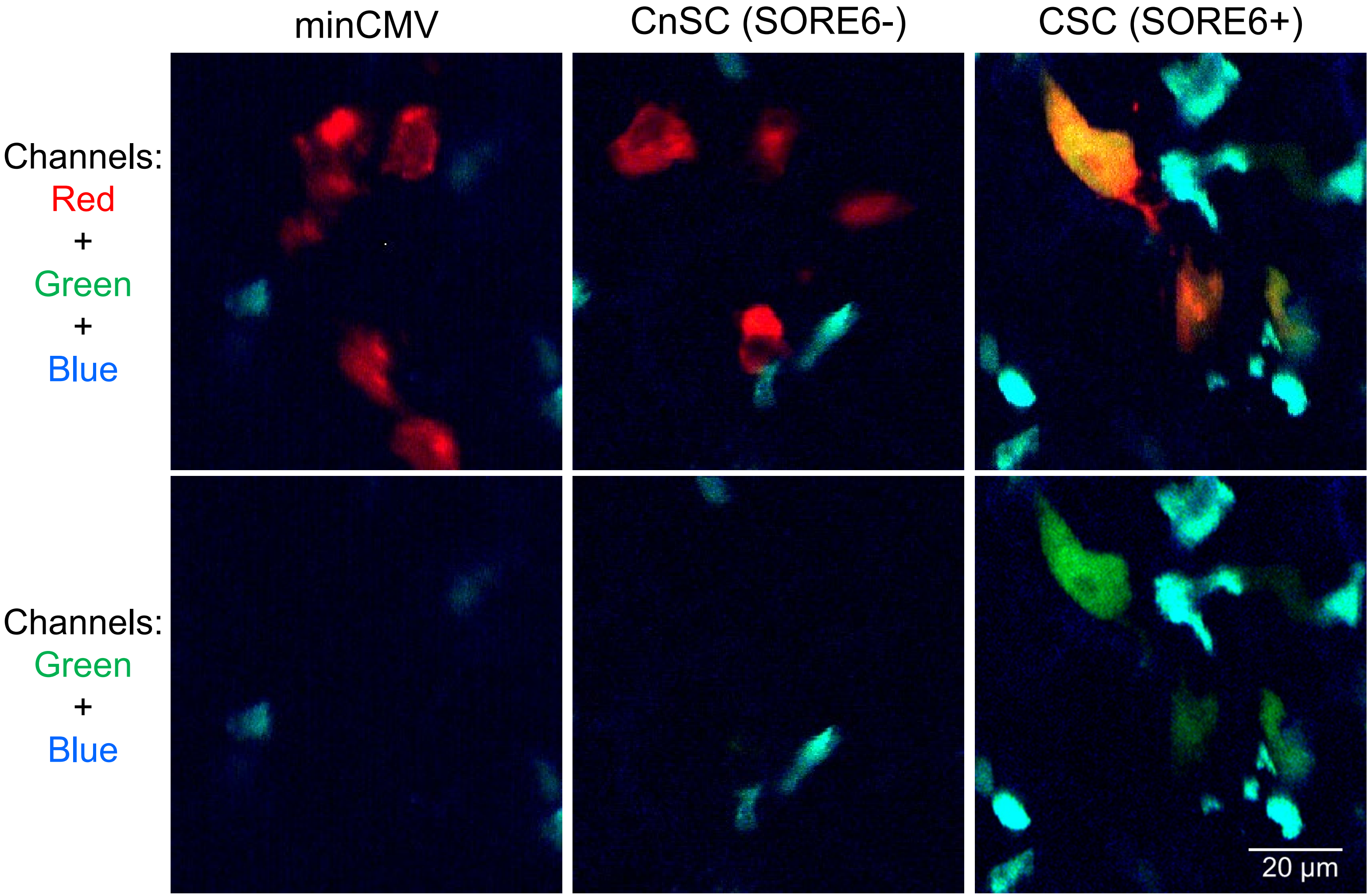


**Supplementary Fig. 4**

**Supplementary Figure 4. Identification of Cancer non-Stem Cells (CnSCs) and Cancer Stem Cells (CSCs) in live lung tissue during intravital imaging.** Panels show representative image of intravital lung imaging: Left: MinCMV control is used to determine the maximum permissible green detector gain setting. Middle: Representative image of tumor cells containing only red fluorescence, identifying them as CnSCs. Right: Representative image of tumor cells containing both red and green fluorescence, identifying them as CSCs. Red = tdTomato and cell tracker dye; Green = GFP fluorescence; Cyan = CFP labeled macrophages.

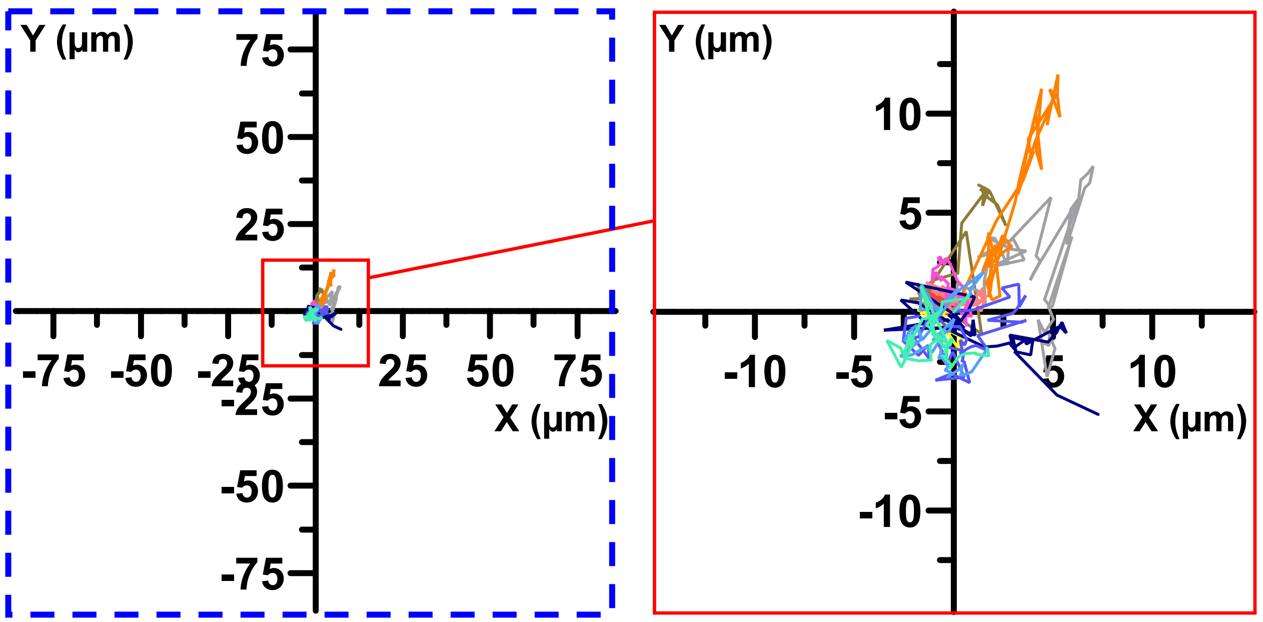


**Supplementary Figure 5. CSCs and CnSCs show minimal motility in the lung.** Blue dashed box indicates the size of an entire field of view. Plot on right is zoom in to the red box in plot on left.

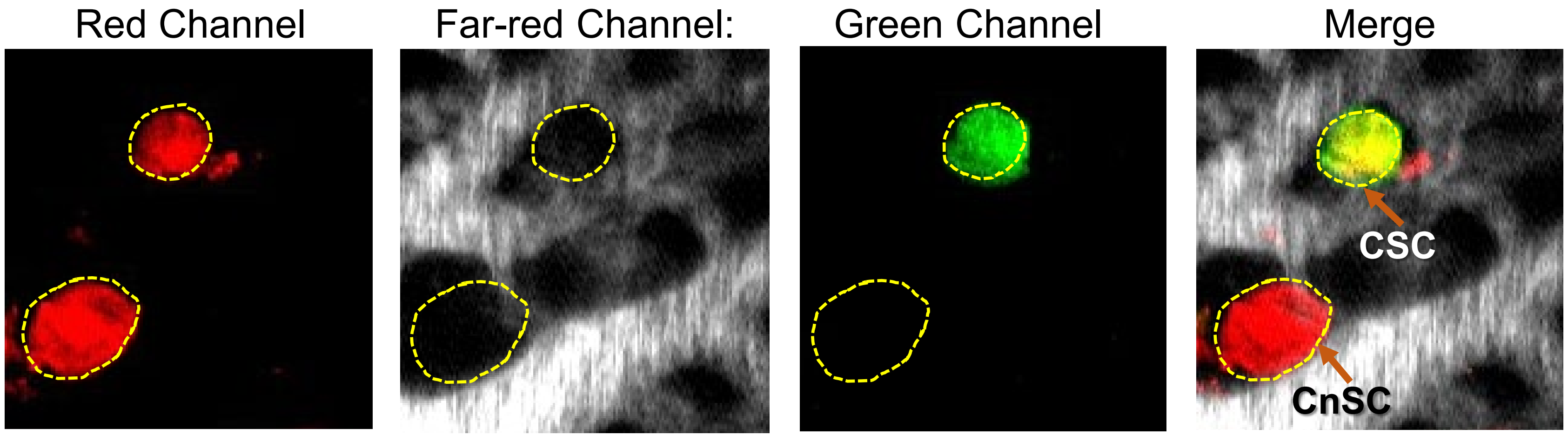


**Supplementary Figure 6. Identification and localization of cancer stem and cancer non-stem cells.** Representative images of tumor cells in the live lung imaged through the lung window. Tumor cells are identified by their red fluorescence (tdTomato and red tracker). Vasculature is labeled with a far-red fluorescent dextran. Cells can be determined to be intra- or extravascular based on their overlap with the far-red signal. Cancer stem cells (CSCs) display green fluorescence from the activated SORE6 reporter. The green (GFP) signal distinguishes GFP-negative CnSCs from GFP-positive CSCs.

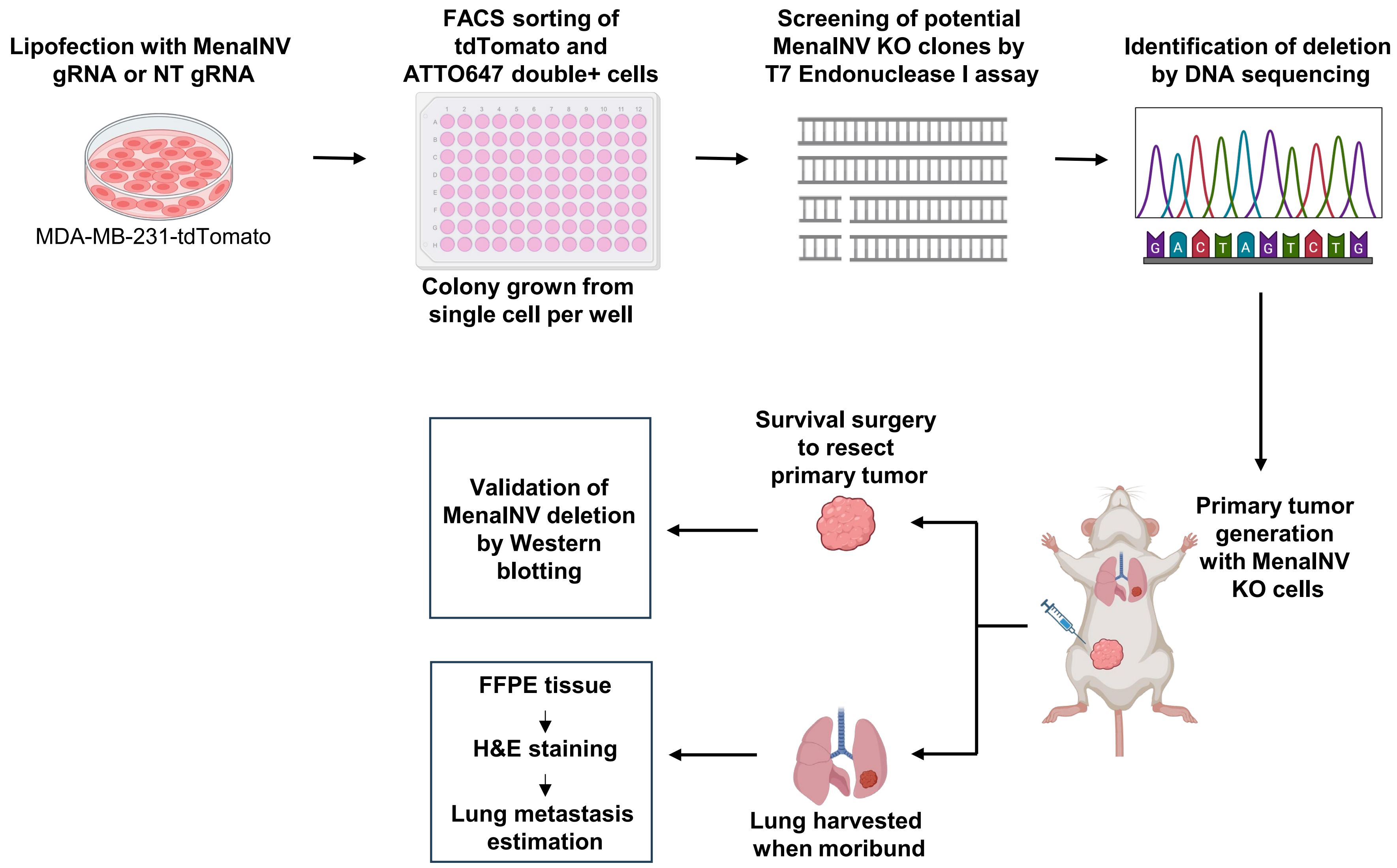


**Supplementary Figure 7. Generation MenaINV knock out MDA-MB-231 cells using CRISPR/Cas9 and validation of MenaINV deletion.** Schematic overview of CRISPR/Cas9 strategy and validation workflow for MDA-MB-231 MenaINV knockout (MenaINV-KO) cells. The image was created with BioRender.com.

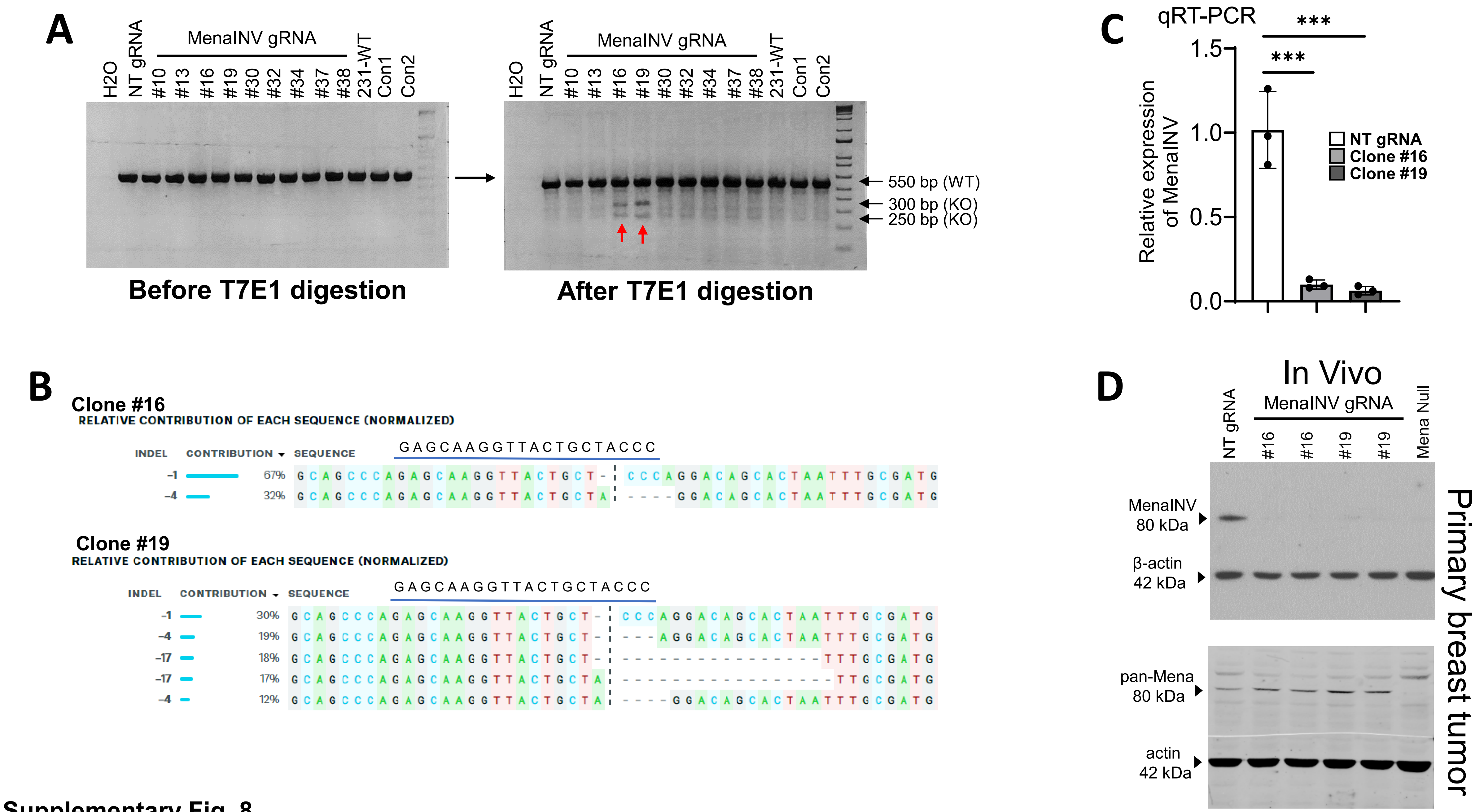


Supplementary Fig. 8

**Supplementary Figure 8. In vitro validation of MenaINV KO clones. A)** PCR amplicons from single-cell derived clones were digested with T7E1 and resolved by gel electrophoresis. Clones #16 and #19 (red arrows) display distinct cleavage bands (~300 bp, ~250 bp) in addition to the uncleaved band (~550 bp), indicating efficient biallelic editing and formation of heteroduplexes with different indels. **B)** Sanger sequencing and ICE analysis for clone #16 show two major indels (-1 bp, 67%; -4 bp, 32%), consistent with a stable biallelic knockout. Clone #19 displays five substantial indels (-1 bp, 30%; -4 bp, 19%; -17bp, 18%; -17 bp, -17%; -4 bp, 12%), indicating allelic heterogeneity likely due to polyploidy or mosaicism. Knockout scores (99 for #16 and 96 for #19) and model fit (0.99 for #16 and 0.96 for #19) confirm efficient editing of both clones. Sequence of MenaINV gRNA shown by blue line and deletion in MenaINV exon is shown by black dashes. **C)** Quantitative PCR confirms MenaINV downregulation in KOs versus controls (mean±SEM, n=3, normalized to GAPDH, log2ddCT; one-way ANOVA, ***p<0.001). **D)** Western blot analysis of MenaINV and pan-Mena in tumors from control, clones #16 and 19, and tumors from MDA-MB-231 Mena null cells generated by CRISPR/Cas9 lacking all Mena isoforms (Mena null).

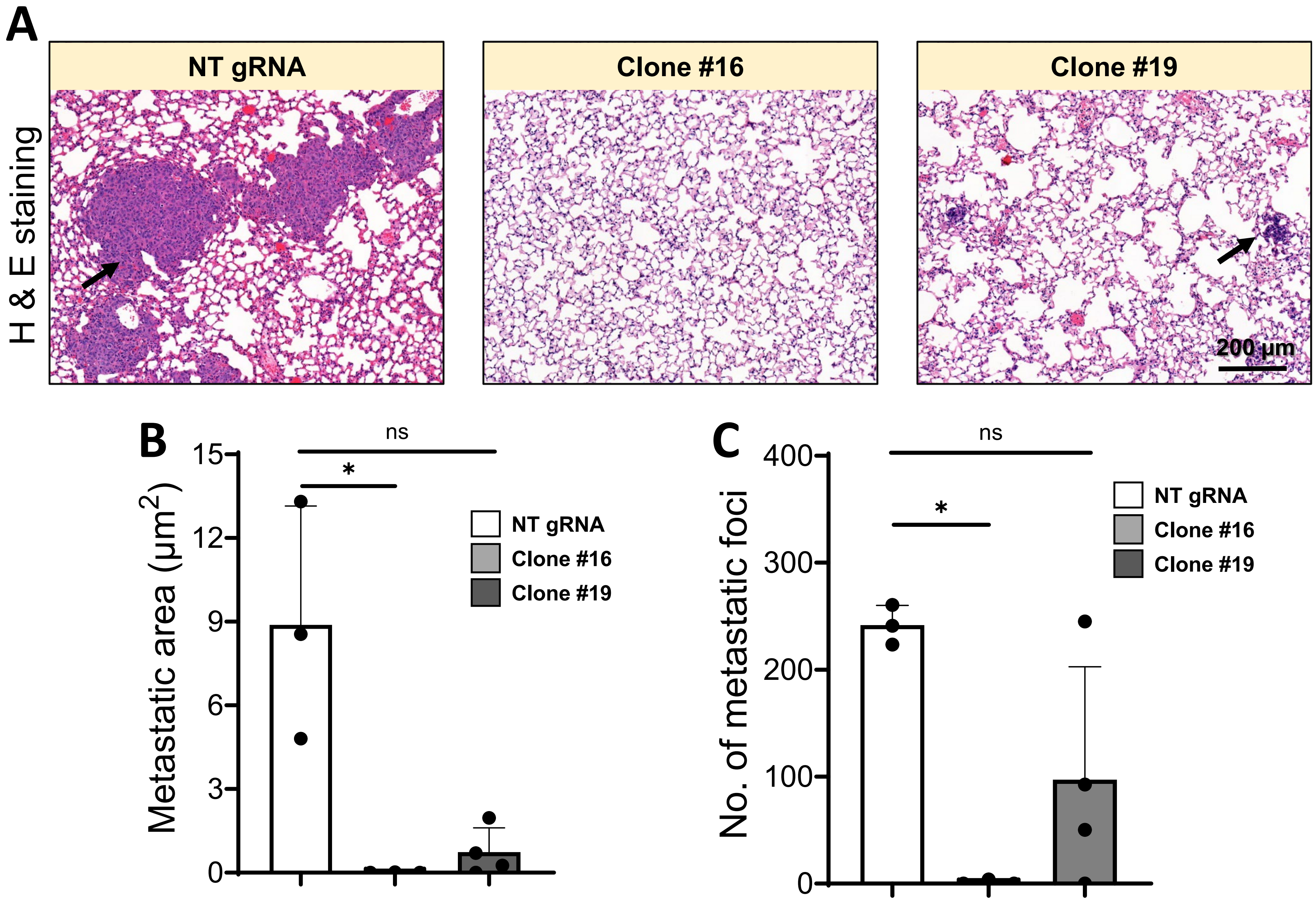


**Supplementary Fig. 9**

**Supplementary Figure 9. Metastases forming ability of MenaINV KO clones**. **A)** Representative images of lungs from mice bearing primary tumors grown from Left: Control (231-tdTomato-NT gRNA), and Middle & Right: two clones of MenaINV-KO (231-tdTomato-MenaINV gRNA) cells showing complete (Clone #16) and partial (Clone #19) abrogation of metastatic capacity compared to Control cells. Metastases are shown by black arrows. **B)** Quantification of metastatic area in H&E-stained lung sections (3 intervals, 50 µm apart, n=3-4/group). Kruskal–Wallis with Dunn's multiple comparison test, *p<0.019). **C)** Quantification of number of metastatic foci (same intervals as in B), n=3-4, Kruskal–Wallis with Dunn's multiple comparison test *p<0.046).

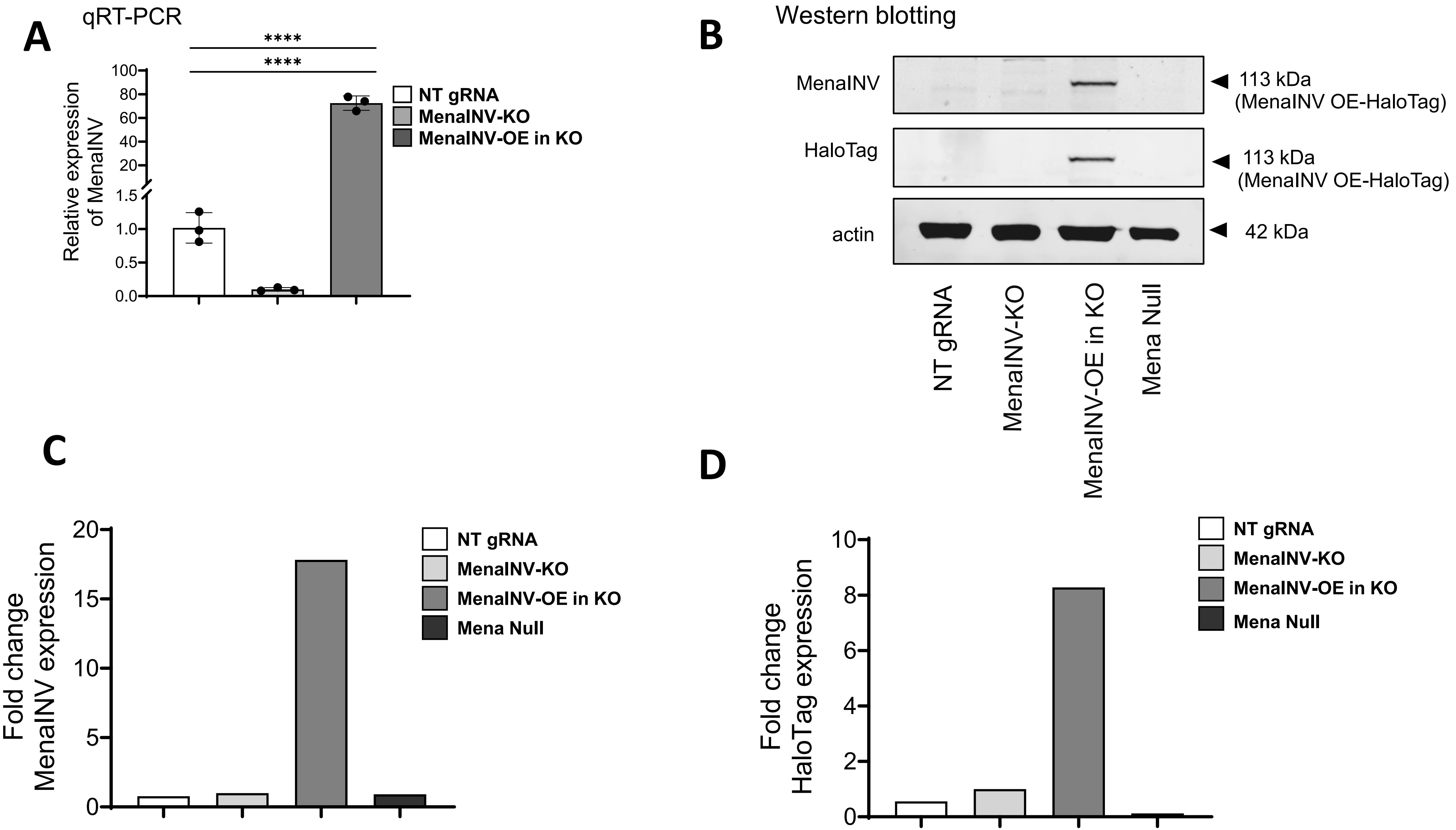


**Supplementary Fig. 10**

**Supplementary Figure 10. Reintroduction of MenaINV expression in MenaINV-KO cells. A)** Quantitative PCR showing successful MenaINV reintroduction in MenaINV-KO cells (mean±SEM, n=3, normalized to GAPDH, log2ddCT; ANOVA, ****$p<0.0001$). **B)** Western blot confirming MenaINV protein reintroduction in MenaINV-KO cells. MDA-MB-231 Mena null cells generated by CRISPR/Cas9 lacking all Mena isoforms identify bands of non-specific antibody staining. **C)** Quantitative RT-PCR demonstrates MenaINV mRNA expression in the overexpression line (MenaINV-OE in KO) is approximately 18-fold higher than wildtype (NT gRNA). **D)** HaloTag protein expression quantification shows approximately 8-fold increase in MenaINV protein levels in MenaINV-OE in KO compared to controls.

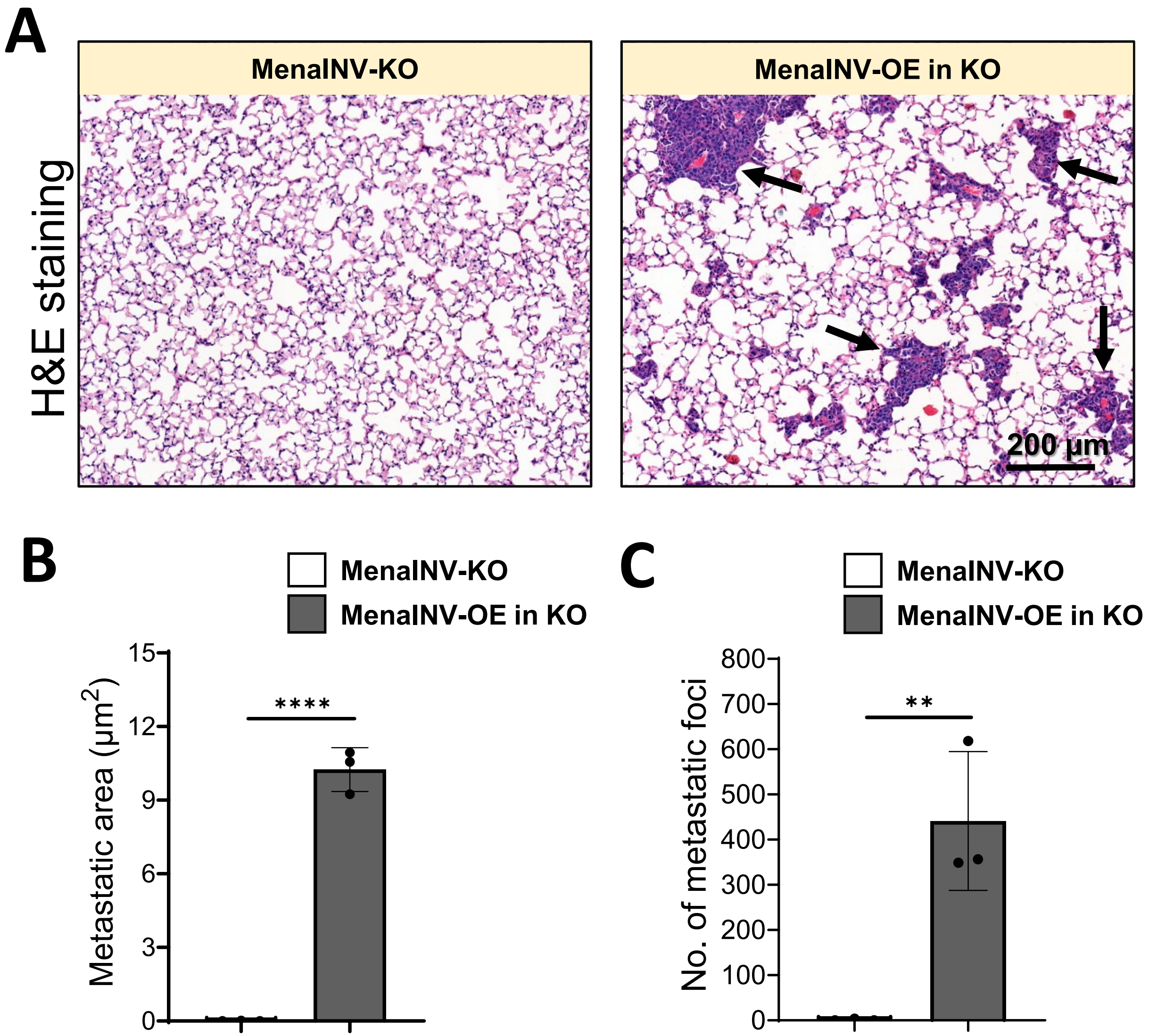


Supplementary Fig. 11

**Supplementary Figure 11. Reintroduction of MenaINV expression in MenaINV-KO cells rescues metastatic potential. A)** Representative images from lungs in spontaneous metastasis model with MenaINV manipulated cells. MenaINV-KO cells result in no metastases, while reintroduction of MenaINV in MenaINV-KO cells (MenaINV-OE in KO) results in the formation of small metastases (black arrows). **B)** Quantification of metastatic area (3 sections, 50 µm apart, n=3/group). Two-tailed Student's t-test, ****$p<0.0001$). **C)** Quantification of metastatic foci (3 sections 50 µm apart, n=3/group). Two-tailed Student's t-test, **$p<0.01$).
